# Invasion and extinction dynamics of mating types under facultative sexual reproduction

**DOI:** 10.1101/632927

**Authors:** Peter Czuppon, George W. A. Constable

## Abstract

In sexually reproducing isogamous species, syngamy between gametes is generally not indiscriminate, but rather restricted to occurring between complementary self-incompatible mating types. A longstanding question regards the evolutionary pressures that control the number of mating types observed in natural populations, which ranges from two to many thousands. Here, we describe a population genetic null model of this reproductive system and derive expressions for the stationary probability distribution of the number of mating types, the establishment probability of a newly arising mating type and the mean time to extinction of a resident type. Our results yield that the average rate of sexual reproduction in a population correlates positively with the expected number of mating types observed. We further show that the low number of mating types predicted in the rare-sex regime is primarily driven by low invasion probabilities of new mating type alleles, with established resident alleles being very stable over long evolutionary periods. Moreover, our model naturally exhibits varying selection strength dependent on the number of present mating types. This results in higher extinction and lower invasion rates for an increasing number of residents.

## 1. Introduction

In isogamous species, the gamete size differentiation that defines the sexes in anisogamous species is absent. Instead, these species produce gametes that are morphologically similar (Lehtonen et al., 2016). Despite this physical similarity, the gametes of isogamous species are typically not interchangeable, but rather fall into one of a number of genetically determined self-incompatible gamete classes, termed mating types. Syngamy can only occur between gametes of distinct and complementary mating types.

The evolutionary explanation for this self-incompatibility, which limits the number of potential mates available to an isogamous organism, has been the subject of debate, with a number of competing hypotheses proposed (Billiard et al., 2011). These include theories that such self-incompatibility alleles limit inbreeding depression (Charlesworth and Charlesworth, 1979), increase encounter rates between gamete pairs (Hoekstra, 1982; Hadjivasiliou et al., 2015; Hadjivasiliou and Pomiankowski, 2019), allow for ploidy level detection and the instigation of the zygote developmental program (Haag, 2007; Perrin, 2012), or manage cytoplasmic conflict by promoting uniparental inheritance of organelles (UPI) (Hurst and Hamilton, 1992; Hadjivasiliou et al., 2012).

Empirical observations of the number of mating types vary between species, ranging from 2 (reminiscent of the two sexes found in anisogamous species) to many thousands (Kothe, 1996). For example, intermediate numbers of mating types are reported in ciliates of the *Tetrahymena*species (3 − 9 different mating types (Doerder et al., 1995; Phadke and Zufall, 2009)) and slime molds (2 − 13 mating types (Bloomfield et al., 2010; Clark and Haskins, 2010)). Larger numbers can be found in fungal populations; in *Coprinellus disseminatus* the global population is estimated to contain 123 different mating types (James et al., 2006), while *Schizophyllum commune* has a staggering 23,328 distinct types (Kothe, 1999). This naturally leads to the question which type of evolutionary pressures govern this diversity in mating type number? This has been the subject of much debate, motivated in part by a discrepancy between these empirical observations and simple evolutionary reasoning.

From a theoretical standpoint, one might naïvely expect to see a very large number of mating types within any given species, due to the “rare sex advantage” of novel types (Iwasa and Sasaki, 1987). Since mating types are self-incompatible, rare types have more opportunities for mating and thus each type experiences negative-frequency dependent selection. Therefore, each novel mating type produced by mutation should establish in the population and the number of mating types should consistently grow. However, this prediction stands in stark contrast to what is observed in the natural world. Although isogamous species with hundreds, or even thousands, of mating types are possible (Kothe, 1996), examples of such species are very rare; the vast majority have very few (typically two) mating types (Hadjivasiliou, 2014). Explaining this discrepancy between theory and empirical observation has been the focus of much work, and multiple theories have been proposed (Billiard et al., 2011).

One prominent hypothesis is that UPI drives the evolution of two mating types (Hadjivasiliou et al., 2012), with larger mating type numbers becoming less stable with the increased complexity of coordinating an organelle donor-receiver program (Hurst and Hamilton, 1992). While recent modelling work has shown that when the frequency-dependent effects of UPI are accounted for, invading UPI does not reduce the expected number of mating types (Hadjivasiliou et al., 2013), perceived empirical support for the theory comes from the fact that many species with more than two mating types have developed mechanisms to ensure homoplasmy without UPI. For instance, in *Paramecium bursaria*, with up to eight mating types (Phadke and Zufall, 2009), and *S. commune*, sexual reproduction is achieved following the exchange of nuclei between cells without cytoplasmic mixing (Birky, 1995). However this in turn leads to an opposing question; if it really is only UPI that limits the number of mating types, why do ciliates and Agaricomycetes, with these alternative methods for ensuring homoplasmy, still feature species with very few types (as low as two and three respectively, see James (2015))?

A further hypothesis (Hoekstra, 1982; Hoekstra et al., 1991), with renewed attention (Hadjivasiliou and Pomiankowski, 2016), suggests that it is cell-cell signalling between gametes that limits the number of mating types. Here co-evolution between two resident types has been shown to potentially limit the evolutionary success of a third mutant mating type. If derived from a resident type, this new mutant must essentially cross a fitness valley before it can develop encounter rates with the residents comparable to their current pairwise encounter rate, limiting its invasion potential (Hadjivasiliou and Pomiankowski, 2016).

In each of these hypotheses, a biologically plausible emergent mechanism is sought that generates a selective advantage to a pair of mating types, thus limiting their number to two. A notable exception can be found in Iwasa and Sasaki (1987). Here it was demonstrated that under certain dynamics for the mating type encounter rate, the advantage to rare mating types could be suppressed. In the limit of infinitely long-lived gametes (that can always survive until a suitable partner becomes available) selection for mating type numbers greater than two could be eliminated entirely. It was verbally suggested that under such a scenario, genetic drift would purge new mating types, limiting their number. This led to a bimodal prediction for the number of types; populations would either have two (given a particular set of encounter rate dynamics and immortal gametes), or have infinitely many otherwise (with each at infinitely low frequency).

While the conditions required for two mating types in Iwasa and Sasaki (1987) were stringent (and indeed, the possibility of intermediate numbers of mating types impossible) it was notable for suggesting that genetic drift may have a key role to play in determining mating type number. In a similar vein, recent work has emphasized the relevance of finite population size null models (i.e. models in which all mating types are phenotypically similar) for addressing the distribution of mating type numbers observed in nature (Constable and Kokko, 2018; Czuppon and Rogers, 2018). These studies stress that even in the absence of species-specific biological processes, the number of mating types in any real finite population cannot be infinite. Instead the expected number of types will arise from balance between mutations (which introduce new mating types) and extinctions (which decrease the number of types), leading to a number of mating types well below the studied population size.

As an explanation for the low number of mating types often observed in isogamous species, this mutation-extinction balance hypothesis may seem at first improbable, particularly in the light of a number of classic population genetic studies of self-incompatibility (SI) alleles in plants (Wright, 1939, 1960, 1964; Ewens, 1964; Nagylaki, 1975; Yokoyama and Nei, 1979; Yokoyama and Hetherington, 1982), see also Clark and Kao (1994) for a review. From a modelling perspective, the dynamics of SI alleles in gametophytic species, those where SI is determined at the haploid pollen stage (Bod’ová et al., 2018), closely resemble those of SI mating type alleles in isogamous species. (A comparison with the dynamics of SI alleles in sporophytic species, in which SI of the haploid pollen is determined by the diploid parent, can be more complicated due to the complex dominance relationships that regulate SI in some of these species (Thompson and Taylor, 1966; Prigoda et al., 2005; Billiard et al., 2007).) In a seminal paper (Wright, 1939), Wright predicted that for the model plant *Oenothera organensis*, a population of around 500 individuals could sustain approximately 13 self-incompatibility alleles (see also Crosby (1966)). As the number of predicted self-incompatibility alleles would rise with increasing effective population size (which intuitively reduces genetic drift and hence extinction rates), many more self-incompatible mating type alleles might naïvely be expected in isogamous species, for which effective population sizes can be of the order 10^6^ (Baranova et al., 2015).

While in the above context the plausibility of the mutation-extinction balance hypothesis might seem doubtful, an important biological feature of isogamous species with less prominence in plants is the potential to reproduce asexually. In many isogamous species, long periods of asexual reproduction are punctuated by rare bouts of facultative sex. For instance, in the single celled green algae, *Chlamydomonas reinhardtii* (with two mating types (Goodenough et al., 2007)), recent genomic estimates have placed the rate of sexual reproduction to be once in every 770 asexual generations (Hasan and Ness, 2018). Sex in yeast, which also typically have two mating types (Butler, 2007), appears to be rarer still, with estimates of once in every thousand to three thousand asexual generations reported in some species (Tsai et al., 2008). In Constable and Kokko (2018) a population genetic model was used to show that for populations within which sex is rare, the relative strength of genetic drift over selection for novel mating types is amplified, leading to a lower expected number of mating types in mutation-extinction balance. This was true even in the absence of any species-specific selective mechanisms. A survey of available empirical data further supported the view that the rate of facultative sex is a key predictor of the number of mating types in isogamous species.

Analytic bounds on the number of expected mating types under a mutation-extinction balance were calculated in Constable and Kokko (2018), using the stationary distribution of a simple Moran-type model of mating type dynamics. In Czuppon and Rogers (2018) an approximation of the expected number of mating types was calculated under the assumption that sex is obligate. Muirhead and Wakeley (2009) used a similar model to calculate the stationary distribution of the frequency spectrum of mating type alleles, however estimates on the number of expected mating types were not explicitly calculated. In addition, facultative sex was not modelled (although the mathematical analysis employed can account for this factor).

While such estimates of the number of mating type alleles in the stationary distribution are informative, they obscure the dynamical processes that drive and maintain this equilibrium, the establishment and extinction of alleles. These quantities are important for two reasons.

Firstly, they are both independent of the mutation rate of new SI alleles. In null models of mating type dynamics the mutation rate is a confounding parameter as no estimate for this rate is available. (A similar problem was encountered by Wright, who conceded that there could be argument about what a “reasonable” mutation rate should be (Wright, 1939)). In contrast, the mutation rate affects neither the establishment probability of a mutant mating type nor the expected extinction time of a resident type. These quantities thus provide mutation rate independent measures of the evolutionary dynamics of mating types, that can be used to test the feasibility of the mutation-extinction hypothesis.

Secondly, whereas the stationary distribution of mating type number can only be observed over evolutionary time periods, the establishment probability of new mating types and the mean extinction time of mating types can be observed over significantly shorter time periods. These quantities are thus important for providing empirically testable insight into the evolution of mating type number.

Extinction rates have previously been studied in related systems featuring negative frequency dependent selection, which results in high levels of polymorphism. In Takahata (1990) a one-dimensional stochastic diffusion was used to identify the dynamics of the polymorphic major histocompatibility complex (MHC) system with a time re-scaled neutral coalescent. This allowed for estimating the extinction rates of MHC alleles. These results were verified numerically in Takahata and Nei (1990) and later in Slatkin and Muirhead (1999). The same theoretical approach was adapted to gametophytic SI in plants in Vekemans and Slatkin (1994) obtaining results showing that the time to the most recent common ancestor of a sex allele might even exceed the speciation time. In the specific context of mating type alleles, such results have thus far been absent. However recently in Czuppon and Rogers (2018) the establishment probability of newly arising mating type was calculated for populations in which sex is obligate. We will rely on this quantity to obtain our estimate on the mean extinction time of resident mating types.

In this paper we complement this literature by exploring scenarios of self-incompatibility under facultative sex. We begin by calculating an analytic expression for the stationary distributions of the number of mating types, extending the results of Constable and Kokko (2018) (where only bounds of the mode of this distribution were calculated). We then provide an expression for the establishment probability of a novel mating type in facultatively sexual populations, generalizing the results of Czuppon and Rogers (2018). Finally, combining these expressions, we calculate the mean time to extinction of a resident mating type allele in a novel way. To be more precise, instead of relying solely on the one dimensional dynamics of a focal mating type (in mating type frequency space) we make use of the dynamics on the number of mating types (in mating type number space). We conclude by discussing the biological implications of these results.

## 2 Model

We consider a population in which the self-incompatible mating type of an individual is determined by one of an infinite set of potential alleles at a single locus. Such a system is for example found in the social amoebae *Dictyostelium discoideum* (with three mating types (Bloomfield et al., 2010)) and *Didymium squamulosum* (with up to 12 mating types (Clark and Haskins, 2010)). The evolutionary dynamics are determined by a Moran process; generations are overlapping and the population is assumed to be at an ecological equilibrium at which the number of individuals, *N*, is constant over time. We denote the number of individuals carrying mating type *i* by *n*_*i*_, such that 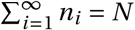.

Asexual reproduction happens with probability *c* ∈ [0, 1]. In this case a randomly chosen individual produces a clone which replaces one of the individuals in the population, also chosen uniformly at random.

During sexual reproduction, which occurs with probability 1 − *c*, two randomly chosen individuals mate if they express different mating types. In case of a successful mating an offspring individual is produced and replaces one of the individuals of the parental population. The mating type of the offspring is chosen at random among the two parental alleles.

Additionally, we consider the emergence of novel mating types through mutations which occur at rate *m*. We implement mutations according to the infinitely many alleles model (Kimura and Crow, 1964), where the mutated individual expresses a completely new mating type not previously present in the population. For simplicity we consider the case in which mutation events are decoupled from reproduction. Mutants possess the same characteristics as any other mating type in the population; that is they are self-incompatible and mate with non-self types at the same per-capita rate as the resident types.

Mathematically implementing the model described above, the probability per unit time for a type *i* to increase by one and a type *j* to decrease by one through a birth-death event is given by

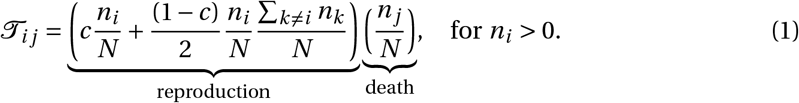

The reproduction term is split into an asexual component (the first term) and a sexual component (the second term). We note, that the sum in the sexual reproduction term goes over all non-*i* mating types present in the population, generating a reproductive advantage to rare mating types. The probability per unit time that a novel mating type *i* is generated from an ancestral mating type *j* is given by

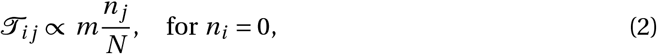

which can only occur when type *i* is not already present in the population. As described in the Supplementary Information, the full expression for the probability transition rate given in Eq. (2) features an additional normalization constant, included to ensure that the mutation rate is independent of the population composition (i.e. that 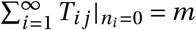).

The deterministic limit (*N* → ∞) of this model is described by a system of ordinary differential equations where the dynamics (neglecting mutations) of a the *i*^*th*^ mating type, with frequency 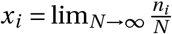, is given by

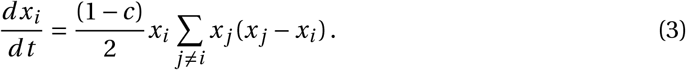

This system has been studied in Iwasa and Sasaki (1987) (see their Mating kinetics I). The dynamical system possesses an internal stable fixed point where all mating types are present at equal frequencies. Analogous finite population size models lead to stochastic differential equations and have been analyzed in Czuppon and Rogers (2018). Note that in that paper mutation was implemented in a distinct manner, with the assumption that mutation occurs <u>with</u> reproductive events. While their implementation is biologically more reasonable, the choice leads to only minor quantitative differences in the dynamics.

## 3 Results

In this section we mathematically analyze the null model just presented. We begin by characterizing the long-time equilibrium behavior of the model, which is described by its stationary distribution. This allows us to answer the key question of how many mating types, *M*, are predicted by the null model as a function of the population size *N*, mutation rate *m* and, importantly, relative rate of asexual reproduction, *c*. However, while this results in an important benchmark for the expected number of mating types in real populations, it provides little insight into the dynamics of mating type number. To this end we further calculate the probability that a novel mutant mating type establishes in the population and the expected extinction time of a resident mating type allele. These quantities provide a deeper insight into the ongoing dynamics of mating type number in real populations.

### 3.1 The stationary distribution of individuals carrying each mating type allele

We denote by 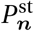 the stationary distribution of the number of mating type alleles of each mating type. In Constable and Kokko (2018) it was shown that an analytic solution for 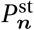 is accessible as the probability transition rates 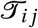 in Eqs. (1-2) can be decomposed into the product of a birth function *b*(*n*_*i*_) and death function *d*(*n*_*j*_) that each depend only on the number of each mating type reproducing and dying respectively;

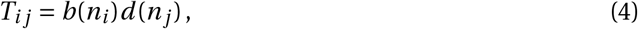

where

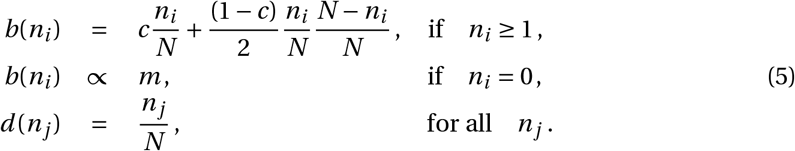

Under this decomposition, the stationary distribution of mating type alleles takes the exact form

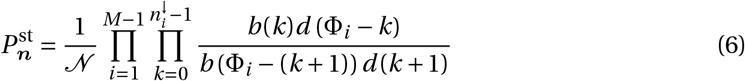

where ***n**^↓^* is the vector *n* reordered with its largest entries first, 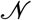 is a normalization constant that enforces 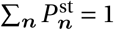 and Φ_*i*_ is defined as

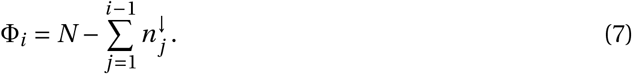

The full derivation of Eq. (6) is given in Constable and Kokko (2018). In terms of birth-death processes, it is interesting to note that Eq. (6) is the product of standard stationary distributions obtained for a two-allele (single variable) population (Karlin and Taylor, 1975), moderated by a series of effective population sizes. A similar observation has been made in a related model of multi-allelic selection (Muirhead and Wakeley, 2009).

### 3.2 The stationary distribution of the number of mating type alleles

We define the stationary distribution of the number of mating types in the population, *M* as 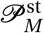. The distribution 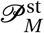 is given by the distribution 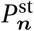 through the following relation;

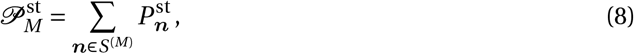

where *S*^(*M*)^ is the set of all vectors ***n*** that represent a population with *M* present mating types (that is, all vectors ***n*** with *M* non-zero elements).

While Eq. (8) can be expressed neatly, evaluating this quantity is problematic as it involves summations of Eq. (6) over sets of the infinite vector ***n***. To make analytic progress we note that when the population size *N* is large and the per-generation mutation rate *m*_*g*_ = *mN* is small, at intermediate times the population resides in a quasi-stationary distribution in the region of the fixed point of the dynamics in the deterministic limit. More precisely, when the population is comprised of *M* mating types, this fixed point is given by *n*_*i*_ ≈ *N*/*M* for each of the *i* present mating types, and *n*_*i*_ = 0 otherwise. Following an extinction or mutation event, thus changing the number of mating types in the population, the population quickly relaxes to a new quasi-stationary distribution in the region of the alternate fixed point with *M* ± 1 mating types, i.e. *n*_*i*_ ≈ *N*/(*M* ± 1). Within the limit of large *N* and small *m*_*g*_ then, 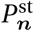 can be approximated by a superposition of these quasi-stationary distributions.

Briefly, the quasi-stationary distribution of the population in the region of a fixed point can be calculated by conducting a diffusion approximation on the underlying Moran model and linearizing the resultant advection-diffusion equation about a deterministic fixed point (the van Kampen approximation). The full calculation is conducted in the Supplementary Information, where we find that the quasi-stationary distribution is normally distributed about the deterministic fixed point with a covariance matrix that can be can be expressed analytically as a function of *N*, *M* and *c* (see Suppĺementary Information Eqn.(S19) and (S20)).

Each of the quasi-stationary distributions can now be renormalized, using Eq. (6) to ‘pin’ the height of each quasi-stationary distribution to the height of Eq. (6) in the region of the deterministic fixed point. The full calculation is detailed in the Supplementary Information, while similar approaches that approximate stationary distributions by a superposition of normal distributions about a stochastic dynamical system’s fixed points can be found in Hufton et al. (2016) and Vasconcelos et al. (2017). Substituting the resulting approximation for 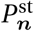 into Eq. (8) and taking the limit of large *N* we find

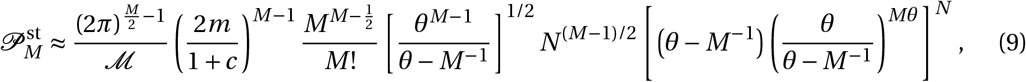

where 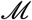 is a normalization constant such that 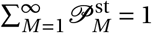 and

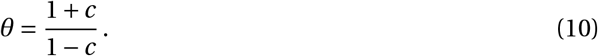

Note that when sexual reproduction is obligate (*c* = 0), *θ* = 1, while when sex is facultative and rare *θ* becomes large. Comparing our approximate expression for the distribution of the number of mating types, Eq. (9), against the results of stochastic simulation of the population we find excellent agreement (see Figure 1).

**Figure 1:**
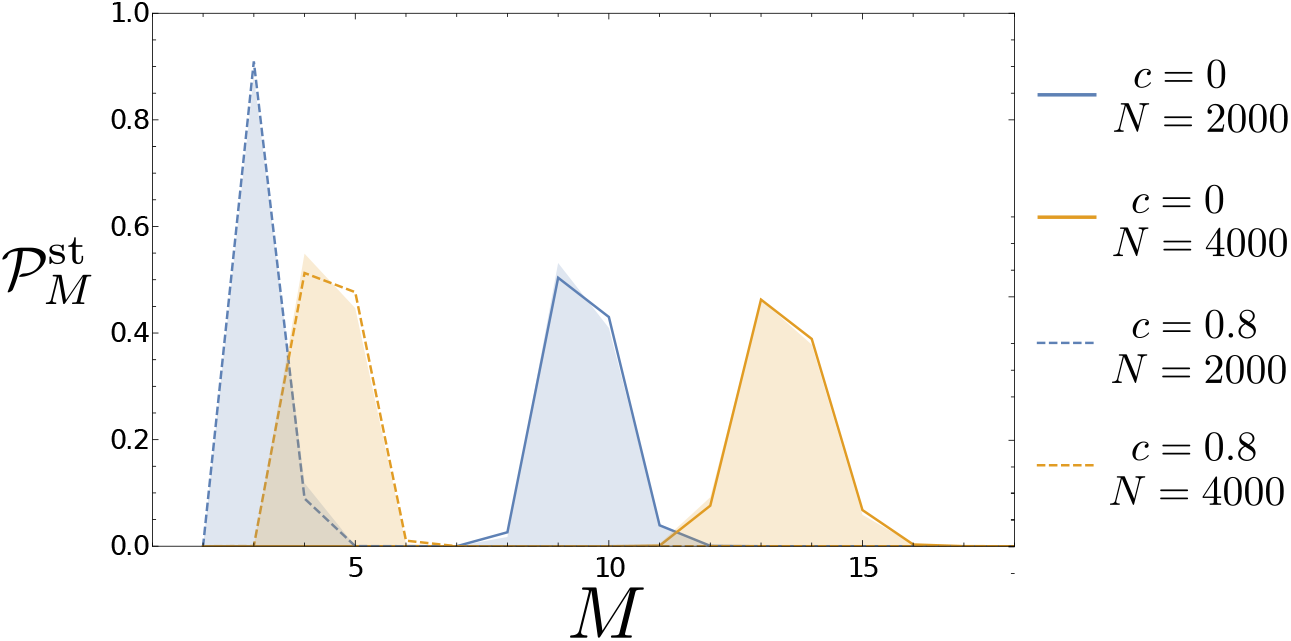
Stationary distribution of mating types, 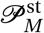. For any given rate of asexual reproduction, *c*, larger populations contain more mating types on average. Meanwhile increasing the rate of asexual to sexual reproduction decreases the expected number of mating types. Analytic results (solid and dashed lines) are obtained by evaluating Eq. (9). Simulation results (corresponding shaded regions) are obtained using the Gillespie algorithm (Gillespie, 1976), averaged over 5 × 10^6^ generations, sampled every 10^2^ generations and following a relaxation period of 10^3^ generations from initial conditions with one more mating type than the mode predicted analytically by Eq. (9). The mutation rate is *m* = 10^−8^ in all plots. The analytic description can be seen to capture the simulated behavior of the model with very high accuracy.

#### 3.2.1 The mode number of mating types

Since the distribution 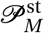 is unimodal, obtaining an estimate for its mode is straightforward numerically. We define by *r*_*M*_ the ratio of probabilities of having *M* and *M* − 1 mating types;

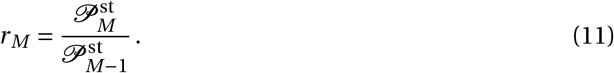

For any given set of parameters, this ratio is independent of the normalization constant 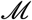. Substituting Eq. (9) into Eq. (11), we obtain a simplified analytic expression for *r*_*M*_ (see Supplementary Information, Eq. (S42)). Since the distribution 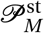 is unimodal, finding its mode is equivalent to finding the value of *M* for which 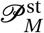 starts to decrease. The approximate mode of 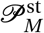, which we denote *M*_o_, can then be obtained as the solution to the equation

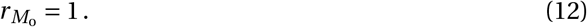

The function *r*_*M*_ − 1 has a single root, and thus solving 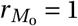 for *M*_o_ is numerically straight-forward.

In Figure 2 we plot the mode number of mating types as a function of the per-generation mutation rate, *m*_*g*_ and the population size, *N*, for four different rates of asexual to sexual reproduction. Here we see that facultative sex has a strong influence over the number of mating types, with even a ratio of 9: 1 asexual to sexual divisions (panel (b)) reducing the expected number of mating types by an order of magnitude compared to the obligately sexual scenario (panel (a)).

**Figure 2:**
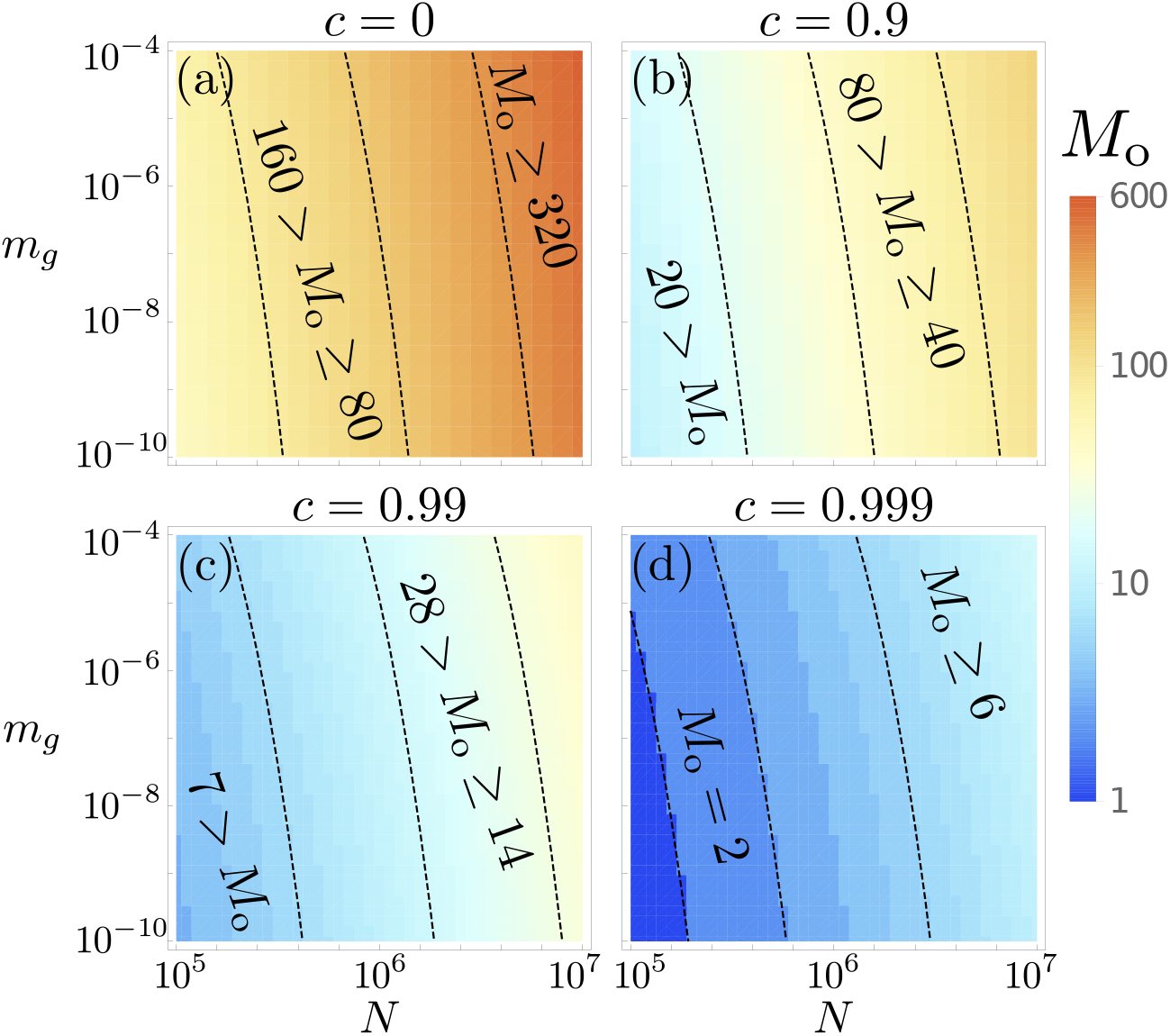
Analytic results on the mode number of mating types as a function of the population size, *N* and the per-generation mutation rate, *m*_*g*_ = *Nm*, under differing rates of asexual reproduction, *c*. As the rate of sexual to asexual reproduction is decreased [panels (a-d)] so too does the expected number of mating types. When sex is obligate [panel (a)], of the order of hundreds of mating types are expected. When sex is facultative and very rare, occurring approximately once to every thousand asexual reproduction events [panel (d)] far fewer mating types are observed. Results are obtained by numerically solving Eq. (12).

### 3.3 The establishment probability of a new mating type allele

Having approximated the mode of the number of mating types we now proceed to study the evolutionary dynamics of the number of mating types. Therefore, we start by computing the probability of a successful establishment, 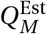, of a newly arising mutant in a population of *M* resident mating types. We define 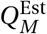 as the probability that this novel mating type allele (initially at a frequency 1/*N*) reaches the new stationary frequency of mating types (1/(*M* + 1)) before any of the resident mating types goes extinct.

For *c* < 1 the deterministic equilibrium is an internal fixed point. This is crucial for our approximation of the establishment probability since we will identify it by the survival probability of a corresponding branching process (see the Supplementary Information). Furthermore, we assume that *N* » *M*, i.e. the population size should be large enough for the resident mating types not to get lost through genetic drift. Under these assumptions we can extend the computation from Czuppon and Rogers (2018) for obligate sexual reproduction and find

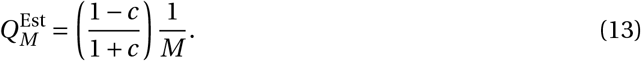

In the case of obligate sex (*c* = 0) the 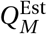 reduces to 1/*M*, the result found in Czuppon and Rogers (2018). For facultative sex (0 < *c* < 1) we see that the establishment probability is reduced. Since asexual reproduction increases the time to reach the stationary state in the deterministic system (i.e. the selection strength for even mating type ratios is reduced, see also Eq. (3)), newly arising mutants spend more time at low frequencies where they are susceptible to be purged by genetic drift. This is reflected by 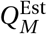 decreasing with *c* (see Figure 3).

**Figure 3:**
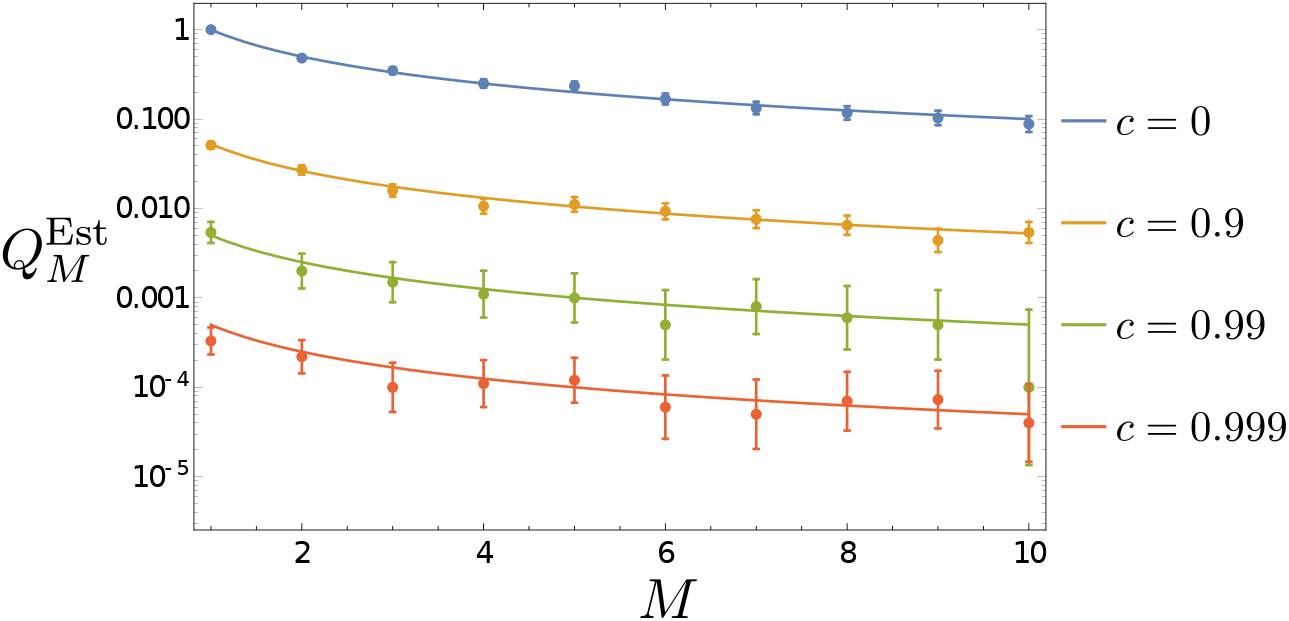
Establishment probability of a novel mating type as a function of *M* for varying rates of asexual reproduction, *c*. Analytic results are obtained by evaluating Eq. (13). Simulation results are obtained from Gillespie simulations; in each plot *N* = 10^5^ and results are averaged over 10^3^ runs (*c* = 0), 10^4^ runs (*c =* 0.9 and *c* = 0.99), and 10^5^ runs (*c* = 0.999). Error bars show Wald-type confidence intervals.

Our definition of the establishment probability includes that none of the resident mating types become extinct during the invasion process. This impedes a straightforward comparison with the case of obligate asexual reproduction, *c* = 1 (i.e. a neutral Moran model). The corresponding establishment probability might naïvely be assumed to be 1/(*M* + 1); however this value ignores the survival of all resident mating types and therefore overestimates the actual establishment probability.

### 3.4 The mean time until the extinction of a mating type allele

Our final goal is to calculate the mean extinction time of a mating type allele. We assume that initially the population is close to its equilibrium (i.e. all mating types are approximately at equal frequencies). We then model the arrival of mating types (through mutation events) and their extinction (due to genetic drift) as a birth-death process on the number of mating types. Let *β*_*M*_ be the probability per unit time that a population with *M* resident mating types gains a new mating type, and *δ*_*M*_ be the probability per unit time that a mating type goes extinct.

Although we are interested in calculating the time between extinction events, we first turn our attention to the stationary probability distribution of this effective ‘birth-death’ process in the number of mating types; 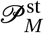. Solving the equations for the stationary distribution of the effective birth-death process, we obtain the well-known result (see for instance (Allen, 2011, Chapter 6))

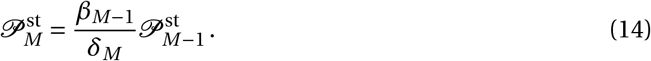

Rearranging for *δ*_*M*_, we find

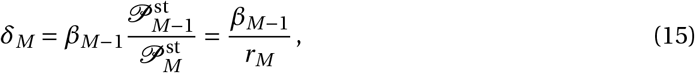

where we have used the definition of *r*_*M*_ given in Eq. (11). The mean time between extinction events is the inverse of this effective death rate, i.e.

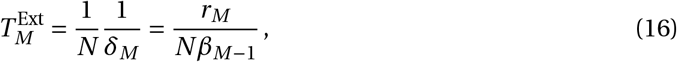

where the factor *N* accounts for the fact that we are measuring the time until an extinction event in time units of generations.

As we have already analytically calculated *r*_*M*_ (see Eq. (11)), all we now need to evaluate the mean time until the extinction of a mating type allele is an expression for *β*_*M*_ (see Eq. (16)). We assume that the effective ‘probability birth rate’, *β*_*M*_ (the probability per unit time that the number of mating types in the system increases) is given by the product of the mutation rate and the establishment probability of a mutant allele (see Eq. (13)):

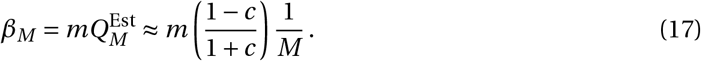

This is an estimate since we argue that invasion happens on a fast time scale when compared to the birth-death chain in *M*−space. However, given very large rates of clonal reproduction (*c* ≈ 1) this assumption may not hold. As *c* increases the deterministic selective forces that drive the mutant towards the stable internal fixed point become weaker. This results in increased establishment times as we have already observed in our analysis of the establishment probability.

Inserting this effective birth rate into Eq. (16) we can express the mean extinction time as

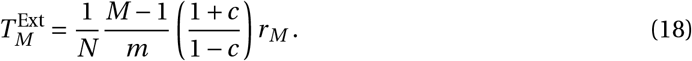

We note that despite the appearance of the mutation rate, *m*, in the above equation, the final expression for 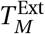 is in fact independent of *m* since *r*_*M*_ is linear in *m*. This can be seen by considering the exponents of *m* in Eq. (11) on the substitution of Eq. (9). Indeed this agrees with our expectations; when mutations are infrequent, the mean time until the extinction of a mating type allele should not depend on the time until a new mating type allele arrives.

The expression derived in Eq. (18) becomes increasingly accurate with increasing population size, *N*, decreasing rates of asexual reproduction, *c*, and smaller numbers of initial mating types, *M*. The approximation can break down however when either *N* is small, or *c* or *M* are large. In this latter range, as the population dynamics become increasingly dominated by genetic drift, various assumptions involved in the derivation of Eq. (18) can become invalid. Most importantly the Gaussian approximation for the quasi-stationary distribution of a focal mating type breaks down. The true distribution becomes increasingly flat, while the Gaussian approximation does not respect the boundary conditions requiring non-negative number of mating types *n*_*i*_. Since this results in erroneous approximations of 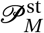, (the crucial ingredient calculating Eq. (14)) when the number of mating types is highly unstable (i.e. when *M* » *M*_*o*_), we do not expect our analytically derived mean extinction time to provide reasonable results in this regime. This reasoning is assessed in more detail and validated in the Supplementary Information (see Section S6).

We now seek a more quantitative measure of when we expect Eq. (18) to remain valid. We first note that naturally 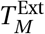 should increase monotonically with the population size, and decrease monotonically with the rate of asexual reproduction and the number of resident mating types. However, on evaluating Eq. (18), we find that these expectations are violated as the relative strength of genetic drift increases (i.e. as the initial number of mating types, *N*/*M*, becomes small) or, conversely, as the strength of selection for equal numbers of mating types decreases (i.e. *c* becomes large). In these regimes, the dynamics of the mating type alleles approach neutrality, and the extinction time is better approximated by standard results on the neutral multi-allelic Moran model Baxter et al. (2007) (see also the Supplementary Information);

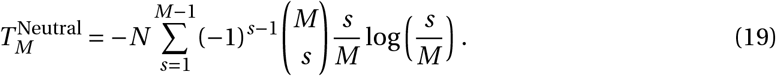

Combining 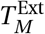 and 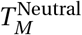, the mean time to extinction of a mating type allele can then be approximated by

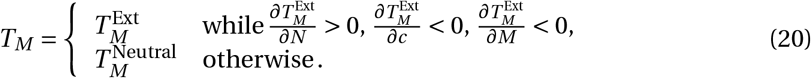

In Figure 4, we can see that this captures the results of simulations very well.

**Figure 4:**
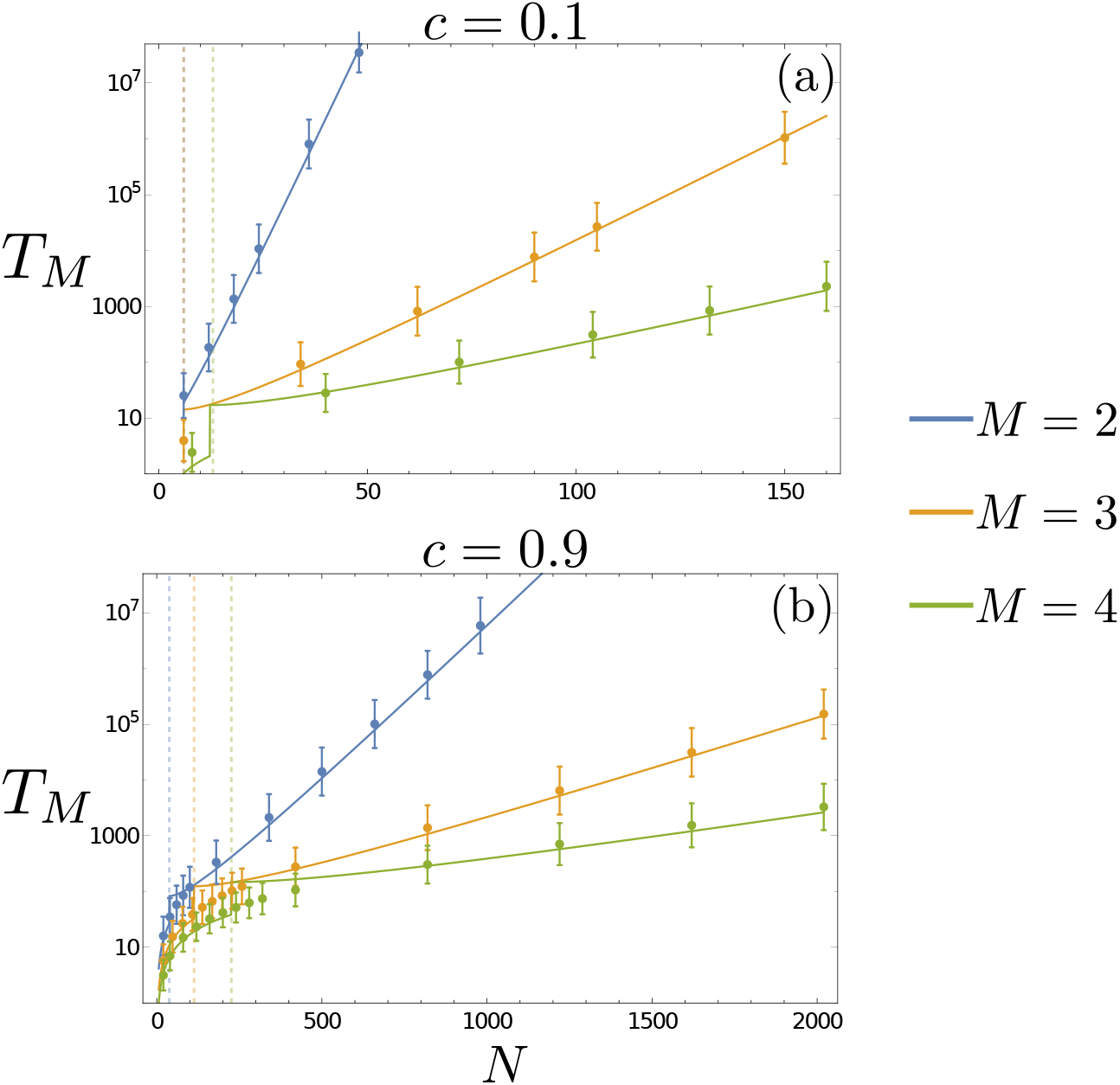
Mean extinction time of the first mating type allele, *T*_*M*_, as a function of *N* for populations with initially *M* = 2, *M* = 3 and *M* = 4 mating types. Analytic results (see Eq. (20)) are plotted as solid lines. Simulation results are averaged over 10^3^ Gillespie simulations, with error bars indicating the standard deviation of results. Vertical dashed lines indicate the transition between the approximation of Eq. (18) and the neutral limit of Eq. (19) (see also Eq. (20)). As the rate of asexual reproduction is increased (from *c* = 0.1 in panel (a) to *c* = 0.9 in panel (*b*)), the mean extinction time drops rapidly (note differing scales on *x*-axis).

In Figure 5 we plot predicted extinction times for regions of parameter space that are prohibitively time consuming to investigate numerically. The gray regions which dominate the parameter space for low asexual reproduction rates depict extinction rates larger than 10^11^ generations. To set this into context, we compare this to the evolutionary history of fungi. Fungi first evolved around 1.5 billion years ago (Wang et al., 1999). Assuming approximately 100 generations a year, a conservative estimate considering the doubling time of yeast which is around 90 minutes, this leads to an order of magnitude guess of 10^11^ generations for the entire evolutionary history of fungi. Hence, in the context of obligate sex (see Figure 5 top-left panel), mating types alleles are expected to remain fixed in a population while their number remains less than 6. This condition is severely relaxed in populations in which sex is facultative (see for example Figure 5 bottom-right panel). Even though extinction times remain high for realistic effective population sizes, some turnover of mating types might be expected over long time periods, e.g. the evolutionary history of fungi. Moreover, this observation indicates that mating types loss is most likely be driven by extreme population bottlenecks decreasing the effective population size *N*.

**Figure 5:**
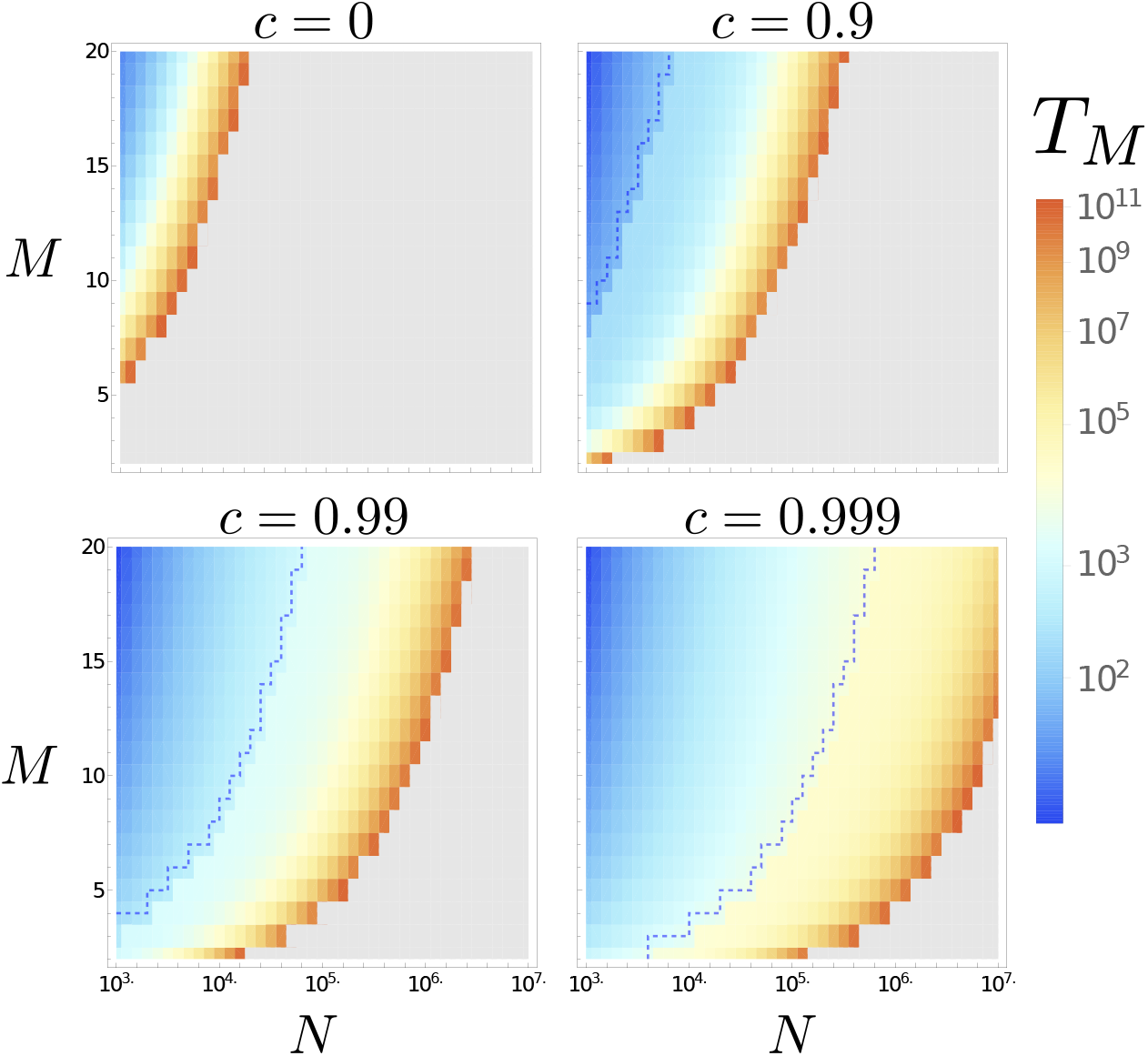
Analytic predictions for the mean extinction time of a resident mating type allele as a function of *N* and *M* (see Eq. (20)). The blue dashed line indicates the parameter regime in which extinction rates become approximately neutral (Eq. (19)). Extinction times in the gray shaded region exceed 10^11^ generations. As these time are approximately longer than the evolutionary history of fungi, they have been omitted for clarity.

## 4 Discussion

The evolutionary mechanisms that drive the number of mating type alleles observed across species has been the topic of numerous theoretical studies. Our analysis in the context of haploid self-incompatibility adds new results and perspectives on the evolutionary dynamics of this type of balancing selection.

A specificity of mating type SI, as studied here, when compared to gametophytic SI, as prevalent in plants, is the possibility to reproduce asexually. Our findings support and extend the previous result that switching between an asexual and a sexual life cycle significantly reduces the number of mating types in a population (Constable and Kokko, 2018). Empirically, available data appears to support the view that more frequent sex is correlated with more mating types (see Table 1 and again Constable and Kokko (2018)). However we are hampered from making any strong empirical claims in this area as a result of paucity of empirical estimates of rates of sex in natural populations. In most species such estimates are absent, however with new methodologies and understanding for estimating the rate of sex arising increasingly frequently (Nieuwenhuis et al., 2018; Ennos and Hu, 2018; Hartfield et al., 2018), it is our hope that this gap in the literature will soon be filled.

**Table 1:** A comparison of empirical values for the number of mating types in various isogamous species with the predictions of the null model. We assume an effective population size of between 10^7^ ≥ *N* ≥ 10^5^ in our calculations of *M* and *T*_*M*_ (note 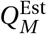 is independent of *N*). In our calculation for *M*, we further assume a per-generation mutation rate, *m*_*g*_ = *mN*, of 10^−6^ ≥ *m*_*g*_ ≥ 10^−8^. For references to empirical data, see the list at the end of the manuscript. For a visualization of the full theoretical distributions of *M*, 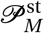, for each parameter set, see Supplementary Information.

| Species | Estimated $c$ | Empirical $M$ | Null model $M_o$ | $Q_M^{\text{Est}}$ | $T_M$ (Generations) |
| --- | --- | --- | --- | --- | --- |
| *S. cerevisiae* * | $1 - 1/2000$ [1] | 2 [5] | $9 \geq M_o \geq 1$ | $Q_2^{\text{Est}} = 1.25 \times 10^{-4}$ | $10^{274} \geq T_2 \geq 10^6$ |
| *C. reinhardtii* | $1 - 1/770$ [2] | 2 [6] | $14 \geq M_o \geq 1$ | $Q_2^{\text{Est}} = 3.25 \times 10^{-4}$ | $10^{707} \geq T_2 \geq 10^9$ |
| Tetrahymena † | $1 - 1/100$ [3] | 3 – 9 [7] | $38 \geq M_o \geq 4$ | $Q_3^{\text{Est}} = 1.68 \times 10^{-3}$<br>$Q_9^{\text{Est}} = 5.58 \times 10^{-4}$ | $10^{1822} \geq T_3 \geq 10^{20}$<br>$10^{153} \geq T_9 \geq 10^4$ |
| *S. commune* | 0 [4] | 23,328 ‡ [8] | $510 \geq M_o \geq 48$ | $Q_{23,328}^{\text{Est}} = 4.29 \times 10^{-5}$<br>$Q_{288}^{\text{Est}} = 3.47 \times 10^{-3}$<br>$Q_{81}^{\text{Est}} = 1.23 \times 10^{-2}$ | $41 \star > T_{23,328} \geq 0.5 \star$<br>$10^{26} \geq T_{288} \geq 58 \star$<br>$10^{337} \geq T_{81} \geq 10^4$ |

\* Considering only strains of *S. cerevisiae* that do not feature mating type switching, for which our model is appropriate.
† Considering only Tetrahymena species with synclonal inheritance, for which our model is appropriate.
‡ Tetrapolar with mating type compatibility determined by genetic complexes *A* (with 288 variants) and *B* (with 81 variants).
★ Lower-bound extinction time is effectively neutral (see Eq. (20)).

In Table 1 we test our model quantitatively against four species where estimates for the rate of sex are available. We compare the number of mating types observed empirically with the mode number predicted theoretically. Since estimates for the mutation rate of new mating type alleles and the effective population size parameters are difficult to obtain, we considered a range of values. We find that while facultative sex explains much of the variation in mating type number, there are quantitative disagreements. In particular, for a range of parameters (particularly large effective population size) the number of mating types is overestimated in *Saccharomyces cerevisiae*, *C. reinhardtii* and Tetrahymena, while it is underestimated in *S. commune*.

Given the simplicity of the model that we have proposed, these quantitative disagreements are not unexpected. An interesting open problem is to explore other null models that are capable of explaining these discrepancies. Essentially this task translates to identifying additional mechanisms that decrease selection for more mating types (where their number is overestimated) or increase this selection strength (where their number is underestimated). There exist a number of biologically reasonable potential candidates.

In Eq. (1) we have assumed mass action encounter rate dynamics between gametes, leading to a linear relationship between a mating type’s frequency and its probability of finding a sexual partner (i.e (1 − *x*_*i*_)). This implementation neglects active mate search. In reality, in species such as *C. reinhardtii* or the diatom *Ditylum brightwellii*, that have developed active methods to increase the encounter rate between complementary mating types (Snell and Goodenough, 2009; Waite and Harrison, 1992), this term might be more accurately described by a decreasing concave up function (see for example Ashby and Gupta (2014)). This would qualitatively recapitulate the results of Iwasa and Sasaki (1987) (Mating Kinetics 3 and 4) and lead to a decrease in the predicted number of mating types.

Further, our model assumes that a mating type is determined by a single locus. Thus, our model is restricted to bipolar mating type systems, as opposed to tetrapolar systems where the mating type is determined by alleles at two loci (Nieuwenhuis et al., 2013). We expect that additional loci may lead to larger numbers of mating types maintained in the population (as mating type combinations can be regenerated through recombination, the extinction rate of mating type alleles will be reduced). Indeed, this would be consistent with the larger number of mating types empirically observed in the tetrapolar *S. commune*, as well as the general observation that tetrapolarity is associated with an increase in mating type diversity (Nieuwenhuis et al., 2013). While it is interesting to note that the number of alleles predicted by our model capture the number of *A* and *B* self-incompatibility complexes in *S. commune* separately (see Table 1), theoretically addressing the mutation-extinction hypothesis for tetrapolar systems in a systematic way, along with the effects of encounter rate dynamics, will be interesting areas for future investigations.

Thus far we have focused on modelling considerations that may well strengthen the mutation-extinction balance hypothesis. Of course, non-neutral differences between mating types must be considered as well, and a pluralistic view that incorporates the remaining hypotheses for observed mating type number may be necessary. For instance, we have not accounted for the fact that newly arising mutants are unlikely to be fully compatible with resident mating types, a biologically important consideration (Power, 1976; Hadjivasiliou and Pomiankowski, 2016) that would lower the number of types relative to our idealized estimate. In addition, we have not considered how forms of homothalism (i.e. the evolution of self-compatibility) may affect our results. For instance most yeasts have evolved the ability to switch mating types between sexual generations (Nieuwenhuis et al., 2018), while among ciliates some species exhibit probabilistic mating type expression (Paixao et al., 2011). These factors make a comparison of our model with such species problematic. Further mechanisms that may limit the number of mating types that are not included in our model, including UPI and costly mate search are reviewed in Billiard et al. (2011).

Our model for haploid SI is conceptually similar to extensively studied gametophytic SI systems observed in plants. Yet we take a novel mathematical approach in estimating the number of mating types supported by a finite population. Instead of using the extinction boundaries of the one-dimensional diffusion approximation of a focal mating type as in most previous studies (Wright, 1939, 1960, 1964; Ewens, 1964; Yokoyama and Nei, 1979; Yokoyama and Hetherington, 1982), we study a birth-death process on the total number of mating types. Utilizing just the local description of the stationary distribution around the interior stable fixed point, we circumvent the problem of the diffusion approximation for systems with a stable deterministic behavior being inaccurate at the boundaries (Assaf and Meerson, 2017). This leads to robust predictions for the stationary probability distribution 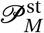 (see Figure 1).

Additionally, we compute the mean extinction time of a mating type allele. Again, we deviate from the approaches taken before which rely on the one-dimensional dynamics of a focal mating type (Takahata, 1990; Vekemans and Slatkin, 1994). This previous method compares the one-dimensional diffusion to a time-rescaled neutral coalescent, thus obtaining estimates for the diversification rate (i.e. the establishment rate of a novel mating type). This comparison is conducted by assuming constant selective strength. In our model, this would be equivalent to assuming that the term Σ_*j*_*x*_*i*_*x*_*j*_ is constant. However, with a varying number of present mating types (and indeed, also with fluctuations in the mating type frequencies themselves) this value is not constant in time. Hence, a comparison with a time-rescaled neutral coalescent does not seem appropriate since the selection strength would need to be re-assessed after each coalescence event which is associated with a loss of a mating type. This variation in selection strength has been empirically observed by analyzing the genome of various *Coprinus cinereus* (mushroom fungus) populations worldwide that show that selective forces in the mating type dynamics are dependent on the number of distinct lineages (May et al., 1999); see also Richman (2000) for other examples in the context of balancing selection. Our technique, explicitly computing the mean extinction time by the effective birth-death process on the number of mating types, avoids this problem. Since the varying selective strength enters in both the establishment probability and the stationary distribution, our computed extinction time accounts for the fluctuating selective pressure.

These explicit estimates on establishment probabilities and typical extinction times yield new insight into the dynamics that drive the low number of mating types predicted by our model in the rare sex regime. One might initially suspect that these numbers are the result of a high turnover of mating type alleles, i.e. frequent extinction and invasion events. However we find that in fact they are a result of very low invasion probabilities for novel mating types, combined with rapidly increasing extinction times as a function of resident mating type diversity. It is worth mentioning that although our approximations for the extinction time break down when the resident state becomes highly unstable, these represent states that would be very short lived in nature, and thus are not relevant from a biological perspective.

Finally, in Table 1 we link our theoretical observation of small numbers of mating type alleles yielding very large extinction times (see Figure 4) to some species examples. Indeed, the large extinction times found by the quantitative study are in line with previous empirical observations of long terminal branches in allelic genealogies under negative-frequency dependent selection, e.g. in fungi (May et al., 1999) and solanaceae (Uyenoyama, 1997). Similar to previous simulation studies (Slatkin and Muirhead, 1999; Gervais et al., 2011) we find that the number of resident mating types strongly influences the diversification rate; the larger the resident number, the lower the diversification rate. This slowdown is ubiquitously observed in natural systems under balancing selection, such as self-incompatibility in fungi (May et al., 1999) or solanaceae (Uyenoyama, 1997) and MHC-systems (Solberg et al., 2008). We find that long terminal branches in self-incompatibility systems, corresponding to old mating type allele ages (with some being older than the corresponding species age (Richman, 2000)) emerge naturally under balancing selection for two reasons: (i) the stability of the internal equilibrium can lead to extremely large extinction times of alleles thus enabling such long branches and (ii) the lower establishment rate due to a large number of SI alleles slows down the diversification rate and hence decreases the number of newly established mating types.

In conclusion, we analyzed the evolutionary dynamics of self-incompatible mating types in facultatively sexual isogamous species. Our results refine previous estimates on the number of mating types maintained in a finite population as well as on the establishment probability of a newly arising mating type allele. Furthermore, we have used these results to compute the mean extinction time of a focal mating type via an effective birth-death process describing the number of mating types. This estimate naturally incorporates variation in selection strength due to varying numbers of resident mating types, a fact that previous studies have failed to incorporate. We are therefore able to qualitatively explain the empirically and numerically observed slowdown of allelic diversification rates in populations under balancing selection. The here presented methodology is theoretically extendable to other systems exhibiting negative-frequency dependent selection.

## Acknowledgements

We thank the Department of Evolutionary Theory at the Max Planck Institute for Evolutionary Biology in Plön for inviting GWAC, resulting in this collaboration. PC received funding from the Agence Nationale de la Recherche, grant number ANR-14-ACHN-0003-01 provided to Florence Débarre. GWAC thanks the Leverhulme Early Career Fellowship provided by the Leverhulme Trust for funding. We also thank Deborah and Brian Charlesworth for their insightful feedback when presenting this work and Sylvain Billiard for directing our attention towards diversification rates of mating types.

## Citations for Table 1

[**1**] As addressed within the discussion, our model is only strictly appropriate for populations in which the mating type does not switch between generations. Mating type switching is prevalent among many Saccharomyces, however most populations feature non-switching strains (Nieuwenhuis et al., 2018). We assume *S. cerevisiae* has a similar ecology to *S. paradoxus*, where sex has been estimated to occur between once every thousand and three thousand generations (Tsai et al., 2008) (indeed, estimates for the rates of outcrossing in both species are comparable).

[**2**] The rate of sex in *C. reinhardtii* has been estimated to once in every 770 generations (Hasan and Ness, 2018).

[**3**] We assume that synclonal Tetrahymena species (such as *T. americanis*, see [7]) feature similar rates of sex to *T. thermophila*. Our estimate on the rate of sex is based on a minimal sexually immature period of 60 − 100 fissions in *T. thermophila* (Doerder et al., 1995). We note however that the frequency of cellular conjugation events observed in other ciliates indicates that lower rates of sex may be appropriate (Lucchesi and Santangelo, 2004).

[**4**] Molecular analysis suggests that *S. commune* features some of the highest rates of sex within the fungal kingdom (Nieuwenhuis and James, 2016). In addition, most of the Agaricomycotina (the class to which *S. commune* belongs) are known to be obligately sexual (Nieuwenhuis and Aanen, 2012). We therefore assume a rate of asexual reproduction of *c =* 0 for *S. commune*. We note however that the life-cycle of *S. commune* features more complicated dynamics than accounted for by our model; it is multicellular and can exhibit vegetative growth as a haploid mycelium.

[**7**] Tetrahymena species *T. americanis*, *T. hegewischi*, *T. hyperangularis* and *T. pigmentosa* have mating type numbers nine, eight, four and three respectively (Phadke and Zufall, 2009). We take care to focus only on these synclonal species, defined as those in which mating type is inherited deterministically based on parental genotype. In caryonidal species, such as *T. thermophila*, the mating type of progeny is determined stochastically (Phadke and Zufall, 2009), and thus a comparison of our model with these species is not appropriate.

[**8**] The mating type of haploid *S. commune* is determined by two complementary pathways, controlled by unlinked genetic complexes *A* and *B*. Each of these regions consists of two weakly recombining loci, leading to a total of four mating type loci. These loci, denoted *Aα*, *Aβ*, *Bα*, and *Bβ*, have respectively 9, 32, 9 and 9 alleles (Stankis et al., 1992). Full compatibility between two mating types is achieved when both the *A* and *B* complexes of the mates are of different specificities; semi-compatibility occurs when the only one of these complexes is of different specificity (Raudaskoski and Kothe, 2019).

## Data availability

The code and simulation outputs can be found on github: https://github.com/gwaconstable/InvExtDynMatTypes.

