## Supplementary Information for "Invasion and extinction dynamics of mating types under facultative sexual reproduction"

#### Contents

**S1 Master equation of the null model** **2**

**S2 Approximation of the quasi-stationary distribution of a focal mating type** **4**

**S3 Approximation of the stationary distribution of the number of mating types, $\mathcal{P}_M^{\rm st}$** **9**

**S4 The mode of the stationary distribution of the number of mating types** **14**

**S5 Establishment probabilities** **15**

**S6 Breakdown of the mean extinction time calculation** **17**

**S7 Neutral mean extinction time** **20**

**S8 The stationary distribution $\mathcal{P}_M^{\rm st}$ for the parameters given in Table 1.** **21**

#### S1 Master equation of the null model

In this section we present the full mathematical description of the model. As discussed in the main text, the function $\mathcal{T}_{ij}$ (see Eq. (1) in the Main Text) gives the probability per unit time that the number of individuals carrying the mating type $i$ allele in the population increases by one and that of type $j$ decreases by one from an initial state $\boldsymbol{n} = (n_1, n_2, \ldots, n_i, \ldots, n_j, \ldots)$, e.g.

$$\mathcal{T}_{ij} = \left( c\frac{n_i}{N} + \frac{(1-c)}{2}\frac{n_i}{N}\frac{\sum_{k\neq i} n_k}{N} \right) \left( \frac{n_j}{N} \right) \ . \tag{S1}$$

For ease of notation, we now introduce the probability transition rate $T(\boldsymbol{n}'|\boldsymbol{n})$, which gives the probability per unit time that a population transitions to a state $\boldsymbol{n}'$ from a state $\boldsymbol{n}$. For our model, the two quantities are related as

$$\begin{aligned} T(\boldsymbol{n}'|\boldsymbol{n}) &= \mathcal{T}_{ij} \quad \text{if} \quad \boldsymbol{n}' = (\ldots, n_i+1, \ldots, n_j-1, \ldots)\,, \\ T(\boldsymbol{n}'|\boldsymbol{n}) &= 0 \quad \text{otherwise}\,. \end{aligned} \tag{S2}$$

The probability of being in a state $\boldsymbol{n}$ at time $t$, $P_{\boldsymbol{n}}(t)$, evolves according to the master equation [Kam07], which can be compactly expressed as

$$\frac{\mathrm{d}P_{\boldsymbol{n}}(t)}{\mathrm{d}t} = \sum_{\boldsymbol{n}'\neq\boldsymbol{n}} \left[ T(\boldsymbol{n}|\boldsymbol{n}')P_{\boldsymbol{n}'}(t) - T(\boldsymbol{n}'|\boldsymbol{n})P_{\boldsymbol{n}}(t) \right] \ . \tag{S3}$$

This equation can be intuitively understood as follows: The probability of being in a state $\boldsymbol{n}$ increases with the probability that the population transitions into state $\boldsymbol{n}$ from a state $\boldsymbol{n}'$ but decreases with the probability that the population was already in state $\boldsymbol{n}$ and transitioned out of it.

For arbitrary initial conditions, it is difficult to solve Eq. (S3), for $P_{\boldsymbol{n}}(t)$. A simpler quantity is the stationary probability distribution $P^{\mathrm{st}}_{\boldsymbol{n}}$, to which the population relaxes on very long timescales. For a time-homogeneous process $P^{\mathrm{st}}_{\boldsymbol{n}}$ is given by the solution to the set of difference equations

$$\sum_{\boldsymbol{n}'\neq\boldsymbol{n}} \left[ T(\boldsymbol{n}|\boldsymbol{n}')P^{\mathrm{st}}_{\boldsymbol{n}'} - T(\boldsymbol{n}'|\boldsymbol{n})P^{\mathrm{st}}_{\boldsymbol{n}} \right] = 0\,. \tag{S4}$$

In [CK18], it was shown that an analytic solution for $P^{\mathrm{st}}_{\boldsymbol{n}}$ was obtainable for the model defined by Eq. (S2) (see also [CK18]: Supplementary Information). This is given in the Main Text, Eq. (6); denoting by $\boldsymbol{n}^{\downarrow}$ the vector $\boldsymbol{n}$ reordered with its largest elements first, the stationary distribution of the population composition can be expressed as

$$P^{\rm st}_{\mathbf{n}} \propto \prod_{i=1}^{M-1} \prod_{k=0}^{n_i^{\downarrow}-1} \frac{b(k)d\left(\Phi_i - k\right)}{b\left(\Phi_i - k - 1\right)d(k+1)} \,, \tag{S5}$$

where $M$ is the number of non-zero elements of $\boldsymbol{n}$ (i.e. the number of mating types in state $\boldsymbol{n}$), and $b(k)$ and $d(k)$ are given by

$$\begin{aligned} b(n_i) &= c\frac{n_i}{N} + \frac{(1-c)}{2}\frac{n_i}{N}\frac{N-n_i}{N}\,, && \text{if} \quad n_i \geq 1\,, \\ b(n_i) &= \frac{m}{M_{\rm max} - M}\,, && \text{if} \quad n_i = 0\,, \\ d(n_j) &= \frac{n_j}{N}\,, && \text{for all} \quad n_j\,, \end{aligned} \tag{S6}$$

(see also, Eq. (4) in the Main Text) and

$$\Phi_i = N - \sum_{j=1}^{i-1} n_j^{\downarrow}\,. \tag{S7}$$

The parameter $M_{\rm max}$ marks the length of the vector $\boldsymbol{n}$ (i.e. $M_{\rm max}$ is the number of distinct possible mating types in the model, which may exceed $N$). The factor $1/(M_{\rm max} - M)$ in the mutation term (which is suppressed in the main text) is an accounting term that ensures that the total population level mutation rate is $m$, i.e.

$$\sum_{i=1}^{M_{\rm max}} b(n_i)\delta(n_i - 0) = m\,, \tag{S8}$$

where $\delta(n_i - 0)$ is the Dirac delta function. As we take the limit $M_{\rm max} \to \infty$ (see Section S3), the appearance of this factor in our transition rates does not alter our results. The validity of Eq. (S5) as a solution can be demonstrated by its direct substitution into Eq. (S4).

### S2 Approximation of the quasi-stationary distribution of a focal mating type

In this section we will calculate an approximate expression for the quasi-stationary distribution of mating type alleles around one of the fixed points (deterministic equilibria). We expect this approximation to be valid on timescales shorter than those of mutation or extinction events.

The approximation that we will use is equivalent to the Linear Noise Approximation (LNA) or a central limit theorem for Markov processes, see [EK86, Kam07]. The procedure is as follows: First we apply a diffusion approximation to Eq. (S3), to obtain a non-linear advection-diffusion equation for the frequency of mating type alleles. Second we will linearize this resultant equation about its fixed point value, to obtain a form of the equation that is amenable to analytical simplifications. Since we are interested in the distribution of mating type alleles close to the fixed point of a focal mating type allele, we will assume both that $m = 0$ and that there are no extinctions (so that $M$ is fixed to its initial value) for the remainder of this section.

We begin by applying the diffusion approximation to Eq. (S3); we introduce variables $x_i = n_i/N$ that measure the frequency of a given mating type allele in the population. The variables $x_i$ are approximately continuous when the population size $N$ is large. We transform into these variables and conduct a Taylor expansion of Eq. (S3) in the small parameter $1/N$. Truncating at next to leading order, we obtain the following non-linear advection-diffusion equation, the Fokker-Planck equation (FPE),

$$\frac{\partial p(\boldsymbol{x},t)}{\partial \tau} = -\sum_{i=1}^{M} \frac{\partial}{\partial x_i} \left[A_i(\boldsymbol{x})p(\boldsymbol{x},t)\right] + \frac{1}{2N} \sum_{i,j=1}^{M} \frac{\partial^2}{\partial x_i \partial x_j} \left[B_{ij}(\boldsymbol{x})p(\boldsymbol{x},t)\right] \,, \quad \text{(S9)}$$

describing the time-evolution of the continuous probability distribution of the variables $\boldsymbol{x}$, $p(\boldsymbol{x},t)$. The expressions for the advection vector $\boldsymbol{A}(\boldsymbol{x})$ and diffusion matrix $B(\boldsymbol{x})$ can be uniquely defined from the underlying probability transition rates Eq. (S1) using well-practiced standard methods [Kam07].

In the current notation we find that the advection vector in Eq. (S9) is given by

$$\begin{aligned} A_i(\boldsymbol{x}) &= \sum_{j\neq i}^{M} \mathcal{T}_{ij} - \mathcal{T}_{ji}\,, \\ &= \sum_{j\neq i}^{M} x_i \left[c + \frac{(1-c)}{2}(1-x_i)\right] x_j - x_j \left[c + \frac{(1-c)}{2}(1-x_j)\right] x_i\,, \\ &= \left(\frac{1-c}{2}\right) \sum_{j=1}^{M} x_i x_j (x_j - x_i)\,. \end{aligned} \tag{S10}$$

As $\dot{\boldsymbol{x}} = \boldsymbol{A}(\boldsymbol{x})$ in the $N \to \infty$ limit (see [MBR14]), this vector essentially gives the deterministic dynamics of the population. Therefore solving $\boldsymbol{A}(\boldsymbol{x}^*) = 0$ allows us to calculate the deterministic fixed point of the dynamics, which we find lies at $x_i^* = 1/M$ for $i = 1, \ldots, M$. Meanwhile the diagonal elements of the diffusion matrix are given by

$$\begin{aligned} B_{ii}(\boldsymbol{x}) &= \sum_{j\neq i}^{M} T_{ij} + T_{ji}\,, \\ &= \sum_{j\neq i}^{M} x_i \left[c + \frac{(1-c)}{2}(1-x_i)\right] x_j + x_j \left[c + \frac{(1-c)}{2}(1-x_j)\right] x_i\,, \\ &= \sum_{j\neq i}^{M} x_i x_j \left[2c + \left(\frac{1-c}{2}\right)(2 - x_i - x_j)\right]\,, \end{aligned} \tag{S11}$$

while the off-diagonal entries are given by

$$\begin{aligned} B_{ij}(\boldsymbol{x}) &= -\left(T_{ij} + T_{ji}\right)\,, \qquad \forall i \neq j\,, \\ &= -\left\{x_i \left[c + \frac{(1-c)}{2}(1-x_i)\right] x_j + x_j \left[c + \frac{(1-c)}{2}(1-x_j)\right] x_i\right\}\,, \\ &= -x_i x_j \left[2c + \left(\frac{1-c}{2}\right)(2 - x_i - x_j)\right]\,. \end{aligned} \tag{S12}$$

With the non-linear FPE for $p(\boldsymbol{x}, t)$ now defined, we proceed to linearize the system about its deterministic fixed point.

As addressed, the deterministic fixed point of the system with $M$ mating types is given by $x_i^* = 1/M$ for $i = 1, \ldots, M$. We assume that the population relaxes to the vicinity of this fixed point before any mutation or extinction events have had time to occur. If $N$ is large, the population will then approach a quasi-stationary distribution around this fixed point on an intermediate timescale. Fluctuations about the fixed point will be of order $1/\sqrt{N}$ due to the central limit theorem for density dependent Markov processes [EK86]. The LNA [Kam07] utilizes this fact by making the change of variables

$$x_i = x_i^* + \frac{1}{\sqrt{N}}\xi_i \tag{S13}$$

to linearize the FPE around the fixed point. Neglecting terms of order $1/N$ or lower, we obtain the following FPE for $\phi(\boldsymbol{\xi}, t)$, the probability distribution of $\boldsymbol{\xi}$ (see [Kam07]; Eq. (6.4) and surrounding discussion):

$$\frac{\partial \phi(\boldsymbol{\xi}, t)}{\partial \tau} = -\sum_{i,j=1}^{M} J_{ij} \frac{\partial}{\partial \xi_i} \left[\xi_j \phi(\boldsymbol{\xi}, t)\right] + \frac{1}{2} \sum_{i,j=1}^{M} B_{ij}(\boldsymbol{x}^*) \frac{\partial^2}{\partial \xi_i \partial \xi_j} \left[\phi(\boldsymbol{\xi}, t)\right] , \tag{S14}$$

where $J$ is the Jacobian matrix of $\boldsymbol{A}(\boldsymbol{x})$,

$$J_{ij} = \left.\frac{\partial A_i}{\partial x_j}\right|_{\boldsymbol{x}=\boldsymbol{x}^*} \tag{S15}$$

As addressed, we are interested in obtaining the stationary distribution of fluctuations about the fixed point (the quasi-stationary distribution of $\boldsymbol{x}$ around $\boldsymbol{x}^*$). Our first step is to obtain the stationary distribution $\phi^{\mathrm{st}}(\boldsymbol{\xi})$ that is the solution to Eq. (S14) at long times;

$$-\sum_{i,j=1}^{M} J_{ij} \frac{\partial}{\partial \xi_i} \left[\xi_j \phi^{\mathrm{st}}(\boldsymbol{\xi}, t)\right] + \frac{1}{2} \sum_{i,j=1}^{M} B_{ij}(\boldsymbol{x}^*) \frac{\partial^2}{\partial \xi_i \partial \xi_j} \left[\phi^{\mathrm{st}}(\boldsymbol{\xi})\right] = 0\,. \tag{S16}$$

Since Eq. (S16) is linear, $\phi^{\mathrm{st}}(\boldsymbol{\xi})$ is normally distributed [Kam07], with mean $\mathbf{0}$ and a covariance matrix, $\Sigma$, that is the solution to the following Lyapunov equation (see [HJ91]);

$$J\Sigma + \Sigma J + B(\boldsymbol{x}^*) = 0\,. \tag{S17}$$

We now must determine the form of the matrices $J$ and $B(\boldsymbol{x}^*)$, which can be calculated from Eqs. (S10-S12). Substituting $x_M = 1 - \sum_{j=1}^{M-1} x_j$ we find that the Jacobian matrix is diagonal with

$$J_{ii} = -\left(\frac{1-c}{2}\right)\frac{1}{M}\,, \quad \forall i\,, \qquad J_{ij} = 0\,, \quad \forall i \neq j\,, \tag{S18}$$

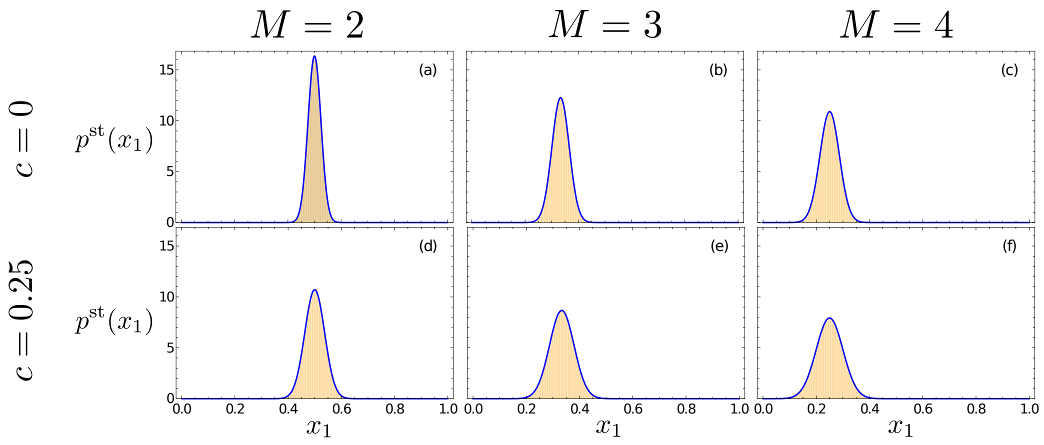


Figure S1: Stochastic simulations (orange histograms) and analytic predictions (blue lines) for the quasi-stationary marginal distribution of $x_1$ in the model. Simulation data is taken from stochastic simulations of a population sampled once every generation for $10^5$ generations. Analytic results are those predicted by the linear Gaussian approximation, Eq. (S25). The population size of $N = 420$ in each figure has been chosen so that the fixed point value $Nx^*$ is integer in each plot. Histogram bin sizes are $1/N$.

while the diffusion matrix evaluated at the fixed point is

$$B_{ii}(\boldsymbol{x}^*) = (1-c)\left(\frac{M-1}{M^2}\right)\left(1-\frac{1}{M}\right) + 2c\frac{1}{M}\left(1-\frac{1}{M}\right), \forall i, \quad \text{(S19)}$$

$$B_{ij}(\boldsymbol{x}^*) = -(1-c)\frac{1}{M^2}\left(1-\frac{1}{M}\right) - 2c\frac{1}{M^2}, \quad \forall i \neq j. \quad \text{(S20)}$$

Since the Jacobian matrix can be expressed by

$$J^{(M)} = -\left(\frac{1-c}{2M}\right)\mathcal{I}, \quad \text{(S21)}$$

where $\mathcal{I}$ is the identity matrix, Eq. (S17) can be simplified to

$$\Sigma = \frac{M}{(1-c)}B(\boldsymbol{x}^*). \quad \text{(S22)}$$

We also note that since the original model is a Moran type model in which the number of individuals $N$ is constant, $J$, $B(\boldsymbol{x}^*)$ and $\Sigma$ are all $(M-1)\times(M-1)$

matrices. Therefore the determinant of $\Sigma$ can be calculated to be

$$|\Sigma| = \frac{1}{M}\left(\frac{1+c}{1-c} - \frac{1}{M}\right)^{M-1} . \tag{S23}$$

This can be verified by solving the matrix of interest, i.e. for a given $M$ with a computer algebra program, e.g. *Mathematica*.

In summary, we find that the stationary distribution of $\boldsymbol{\xi}$ is given by

$$\phi^{\text{st}}(\boldsymbol{\xi}) = \mathcal{N}(\mathbf{0}, \Sigma) , \tag{S24}$$

where $\mathbf{0}$ is the zero vector and $\Sigma$ is given by Eq. (S22). Re-expressing this distribution in terms of our $\boldsymbol{x}$ variables we find

$$p^{\text{st}}(\boldsymbol{x}) \approx \mathcal{N}(\boldsymbol{x}^*, \Sigma/N) , \tag{S25}$$

at intermediate times in the region of the fixed point, as illustrated in Figure S1. Here we see that as the number of mating types in the population, $M$, increases, the mean of the quasi-stationary PDF, $\boldsymbol{x}^* = 1/M$, approaches the extinction boundaries. Simultaneously, increased $M$ leads to increased variance in the quasi-stationary PDF (see Eqs. (S19) and (S22)), though the co-variance of the distribution decreased (fluctuations in each mating type frequency become less correlated, see Eqs. (S20) and (S22)). The variance of the PDF also increases as the rate of asexual to sexual reproduction increases, and fluctuations around the fixed point frequency increase in magnitude (see Eqs. (S19) and (S22)).

Our final step is to re-express Eq. (S25) in terms of the number of alleles of each type in the population. Let $\boldsymbol{\eta}^{(M)}$ be a vector giving an approximate, potentially non-discrete, value of $\boldsymbol{n}$ in the region of a deterministic fixed point with $M$ mating types, e.g.

$$\begin{aligned} \boldsymbol{\eta}^{(M)} &= N\boldsymbol{x}^* , \\ &= \left( \overbrace{\frac{N}{M}, \frac{N}{M}, \ldots, \frac{N}{M}}^{M \text{ elements}}, 0, 0, \ldots \right) . \end{aligned} \tag{S26}$$

For clarity, we choose to write this quasi-stationary distribution as

$$P^{\text{qst}(M)}(\boldsymbol{n}) = \mathcal{N}(\boldsymbol{\eta}^{(M)}, N\Sigma^{(M)}) , \tag{S27}$$

where the superscript $(M)$ emphasizes that the form of this quasi-stationary distribution and the covariance matrix change with the number of mating types present at the fixed point.

### S3 Approximation of the stationary distribution of the number of mating types, $\mathcal{P}_M^{\rm st}$

In the previous two sections, we provided a solution for $P_{\mathbf{n}}^{\rm st}$, the stationary distribution of the population composition, and for $P^{{\rm qst}(M)}(\boldsymbol{n})$, the quasi-stationary distribution of a focal mating type. We will need both of these solutions to obtain an expression for the stationary distribution of the *number* of mating types, $\mathcal{P}_M^{\rm st}$ - our actual object of interest. These stationary distributions of the population composition and the stationary distribution of the number of mating types are related by

$$\mathcal{P}_M^{\rm st} = \sum_{\mathbf{n}\in S^{(M)}} P_{\mathbf{n}}^{\rm st}\,, \tag{S28}$$

where $S^{(M)}$ is the set of all vectors $\boldsymbol{n}$ with $M$ non-zero elements. In other words, to obtain $\mathcal{P}_M^{\rm st}$ we need to sum our expression for $P_{\mathbf{n}}^{\rm st}$ over all the states that contain $M$ mating types. As this calculation is cumbersome, we seek an analytic approximation.

We consider the biologically reasonable parameter regime in which the population size, $N$, is large and the per-generation mutation rate, $m_g = mN$, is small. Under these conditions the population quickly relaxes to a distribution in the region of a deterministic fixed point following an extinction or invasion event. We will use the expression in Eq. (S27) to approximate this quasi-stationary distribution in the region of the fixed point. By doing so, we can replace the sum in Eq. (S28) by a sum of Gaussian distributions (see Figure S2).

The calculation used to obtain Eq. (S27) is based on a linearization of the population dynamics around a deterministic fixed point. As such the expression contains no information about how much more likely the population is to be in the region of a given fixed point with $M$ mating types as opposed to in the region of a fixed point $M+1$ mating types. In order to renormalize each of the normal distributions by the appropriate amount, we "pin" the peak of each normal distribution (see Eq. (S27)) to the height of the full distribution, $P_{\boldsymbol{n}}^{\rm st}$ from Eq. (S5), evaluated in the region of the relevant fixed point. Therefore, we renormalize the height of the Gaussian distribution Eq. (S27) to the height of $P_{\boldsymbol{n}}^{\rm st}$ at $\boldsymbol{n} = \boldsymbol{\eta}^{(M)}$ (see Eq. (S26)). Thus, the probability distribution in the region of a deterministic fixed point with $M$

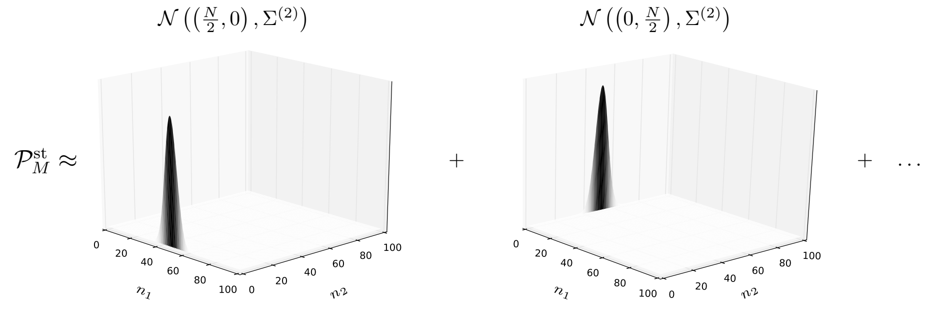


Figure S2: Figure illustrating the analytic approximation used to estimate the distribution of $P_{\boldsymbol{n}}^{\mathrm{st}}$ and thus simplify the calculation of $\mathcal{P}_M^{\mathrm{st}}$ via Eq. (S28).

mating types can be described by the following function

$$P_{\boldsymbol{\eta}^{(M)}}^{\mathrm{st}} \exp\left[-\frac{1}{2N}\left(\boldsymbol{n}-\boldsymbol{\eta}^{(M)}\right)^T\left[\Sigma^{(M)}\right]^{-1}\left(\boldsymbol{n}-\boldsymbol{\eta}^{(M)}\right)\right], \tag{S29}$$

(see Eq. (S27)). Now, the probability of being in the region of a specific fixed point with $M$ mating types is simply the above function integrated over all $\boldsymbol{n}$:

$$\begin{aligned}
P_{\boldsymbol{\eta}^{(M)}}^{\mathrm{st}} & \int_{-\infty}^{\infty}\cdots\int_{-\infty}^{\infty} \exp\left[-\frac{1}{2N}\left(\boldsymbol{n}-\boldsymbol{\eta}^{(M)}\right)^T\left[\Sigma^{(M)}\right]^{-1}\left(\boldsymbol{n}-\boldsymbol{\eta}^{(M)}\right)\right] \\
& \qquad\qquad\qquad\qquad\qquad\qquad\qquad\qquad \mathrm{d}n_1\dots\mathrm{d}n_{M-1} \\
= \quad & P_{\boldsymbol{\eta}^{(M)}}^{\mathrm{st}}\sqrt{(2\pi N)^{M-1}\left|\left[\Sigma^{(M)}\right]\right|} \\
= \quad & P_{\boldsymbol{\eta}^{(M)}}^{\mathrm{st}}\sqrt{(2\pi N)^{M-1}\frac{1}{M}\left(\frac{1+c}{1-c}-\frac{1}{M}\right)^{M-1}} \\
= \quad & \frac{P_{\boldsymbol{\eta}^{(M)}}^{\mathrm{st}}}{\sqrt{M}}\left[2\pi N\left(\frac{1+c}{1-c}-\frac{1}{M}\right)\right]^{(M-1)/2} .
\end{aligned} \tag{S30}$$

Eq. (S30) approximates the probability of being in the region of a particular fixed point with $M$ mating types. The probability of the population having $M$ mating types is therefore given by Eq. (S30) multiplied by the number of fixed points with $M$ mating types. Recall that we introduced $M_{\max}$ as a temporary parameter capturing the length of the vector $\boldsymbol{n}$ (we will shortly take the limit $M_{\max} \to \infty$). The number of ways of choosing a fixed point containing $M$ distinct present mating types from $M_{\max}$ potential mating types is then given by the Binomial coefficient, such that

$$\begin{aligned}\mathcal{P}_M^{\text{st}} &= \sum_{\mathbf{n}\in S^{(M)}} P_{\mathbf{n}}^{\text{st}} \\ &\approx \binom{M_{\max}}{M}\frac{P_{\boldsymbol{\eta}^{(M)}}^{\text{st}}}{\sqrt{M}}\left[2\pi N\left(\frac{1+c}{1-c}-\frac{1}{M}\right)\right]^{(M-1)/2} . \end{aligned} \tag{S31}$$

We now proceed to calculate $P_{\boldsymbol{\eta}^{(M)}}^{\text{st}}$ explicitly.

By substitution of the functions $b(k)$ and $d(k)$ into Eq. (S5), we find after some algebraic simplification [see [CK18]: Eq. (S35)] that

$$\begin{aligned}P_{\boldsymbol{n}}^{\text{st}} \;\propto\; & \left\{\left(\frac{2mN}{M_{\max}-M}\right)\frac{\left[\left(\frac{1+c}{1-c}\right)N\right]!}{(1+c)}\right\}^{(M-1)}\left[2\left(\frac{c}{1-c}\right)N\right]!\,\times \\ & \prod_{i=1}^{M}\left\{\frac{1}{n_i^{\downarrow}}\right\}\left\{\frac{1}{\left[\left(\frac{1+c}{1-c}\right)N-n_i^{\downarrow}\right]!}\right\} .\end{aligned}$$

It is therefore straightforward to show that

$$P_{\boldsymbol{\eta}^{(M)}}^{\text{st}} \;\propto\; \left(\frac{1}{M_{\max}-M}\right)^{M-1}\Omega_M \tag{S32}$$

where

$$\begin{aligned}\Omega_M \;=\; & \left\{\frac{2mN}{1+c}\left[\left(\frac{1+c}{1-c}\right)N\right]!\right\}^{(M-1)}\left[2\left(\frac{c}{1-c}\right)N\right]!\,\times \\ & \left(\frac{M}{N}\right)^{M}\left\{\left[N\left(\frac{1+c}{1-c}-\frac{1}{M}\right)\right]!\right\}^{-M} .\end{aligned}$$

Upon substitution of Eq. (S32) into Eq. (S31), we find

$$\mathcal{P}_M^{\text{st}} \propto \binom{M_{\max}}{M}\left(\frac{1}{M_{\max}-M}\right)^{M-1}\frac{\Omega_M}{\sqrt{M}}\left[2\pi N\left(\frac{1+c}{1-c}-\frac{1}{M}\right)\right]^{(M-1)/2} \tag{S33}$$

As $\mathcal{P}_M^{\text{st}}$ is proportional to the function on the right hand side of Eq. (S33), we can divide through this function by a constant. We choose the binomial coefficient $\binom{M_{\max}}{1}$. The terms involving $M_{\max}$ in Eq. (S33) can then be considered separately, and the limit $M_{\max} \to \infty$ taken;

$$\binom{M_{\max}}{1}^{-1} \binom{M_{\max}}{M} \left( \frac{1}{M_{\max} - M} \right)^{M-1} \underset{M_{\max}\to\infty}{=} \frac{1}{M!}\,. \tag{S34}$$

Thus, for $M_{\max} \to \infty$, our final expression for the approximate stationary distribution of the number of mating types is

$$\begin{aligned} \mathcal{P}_M^{\text{st}} &= \frac{1}{\mathcal{M}} \frac{1}{\sqrt{M}} \left[ 2\pi N \left( \frac{1+c}{1-c} - \frac{1}{M} \right) \right]^{(M-1)/2} \left( \frac{M}{N} \right)^M \left( \frac{2mN}{1+c} \right)^{M-1} \\ &\quad \times \frac{1}{M!} \left\{ \left[ N \left( \frac{1+c}{1-c} \right) \right]! \right\}^{M-1} \left\{ \left[ N \left( \frac{1+c}{1-c} - \frac{1}{M} \right) \right]! \right\}^{-M} \end{aligned} \tag{S35}$$

where we have also absorbed any constant terms that do not involve $M$ (e.g. $[2cN/(1-c)]!$) into the normalization factor $\mathcal{M}$, which is defined so that

$$\sum_{M=1}^{N} \mathcal{P}_M^{\text{st}} = 1\,. \tag{S36}$$

Eq. (S35) can be further simplified by noting that if $N$ is large the terms in the final two factorials are large when $M \geq 2$. We may then express these factorials using the Stirling approximation [AS65];

$$\begin{aligned} \left[ N \left( \frac{1+c}{1-c} \right) \right]! &\approx \left[ 2\pi N \left( \frac{1+c}{1-c} \right) \right]^{1/2} \left[ \frac{N}{e} \left( \frac{1+c}{1-c} \right) \right]^{N(1+c)/(1-c)}, \\ \left[ N \left( \frac{1+c}{1-c} - \frac{1}{M} \right) \right]! &\approx \left[ 2\pi N \left( \frac{1+c}{1-c} - \frac{1}{M} \right) \right]^{1/2} \times \\ &\qquad \left[ \frac{N}{e} \left( \frac{1+c}{1-c} - \frac{1}{M} \right) \right]^{N\left(\frac{1+c}{1-c} - \frac{1}{M}\right)}. \end{aligned}$$

Substituting these expressions into Eq. (S35), we find after some algebra that for $M \geq 2$,

$$\begin{aligned} \mathcal{P}_M^{\text{st}} &\approx \frac{(2\pi)^{\frac{M}{2}-1}}{\mathcal{M}} \left( \frac{2m}{1+c} \right)^{M-1} \frac{M^{M-\frac{1}{2}}}{M!} \left[ \frac{\left(\frac{1+c}{1-c}\right)^{M-1}}{\frac{1+c}{1-c} - \frac{1}{M}} \right]^{1/2} \times \\ &\qquad N^{(M-1)/2} \left[ \left( \frac{1+c}{1-c} - \frac{1}{M} \right) \left( \frac{\frac{1+c}{1-c}}{\frac{1+c}{1-c} - \frac{1}{M}} \right)^{M\frac{1+c}{1-c}} \right]^{N}, \end{aligned} \tag{S37}$$

where we have again absorbed any terms independent of $M$ into the normalization factor $\mathcal{M}$. We can see that Eq. (S37) matches the full expression in Eq. (S35) very well (see also Figure 1 in the Main Text).

#### S4 The mode of the stationary distribution of the number of mating types

Since the distribution $\mathcal{P}_M^{\text{st}}$ is unimodal (see Figure 1, Main Text), determining the mode of $\mathcal{P}_M^{\text{st}}$, amounts to obtaining the first value of $M$ for which

$$\mathcal{P}_{M+1}^{\text{st}} < \mathcal{P}_M^{\text{st}} \,. \tag{S38}$$

This can be obtained with a simple numerical algorithm. Alternatively, we can assume that $M$ is approximately continuous and solve

$$\mathcal{P}_{M+1}^{\text{st}} = \mathcal{P}_M^{\text{st}} \,. \tag{S39}$$

We introduce $r_M$ as

$$r_M = \frac{\mathcal{P}_M^{\text{st}}}{\mathcal{P}_{M-1}^{\text{st}}} \,. \tag{S40}$$

Solving Eq. (S39) is equivalent to finding the root of the following expression

$$r_M - 1 = 0 \,. \tag{S41}$$

To this end we seek to simplify our expression for $r_M$.

Note that as $r_M$ involves the ratio of terms in $\mathcal{P}_M^{\text{st}}$, the normalization factor $\mathcal{M}$ cancels. We obtain

$$\begin{aligned} r_M \;=\; & \sqrt{2\pi}\frac{2m}{1+c}\left(\frac{1+c}{1-c}\right)^{N\left(\frac{1+c}{1-c}\right)+\frac{1}{2}}\left(\frac{M-1}{M}\right)^{\frac{3}{2}-M}\left(\frac{\frac{1+c}{1-c}-\frac{1}{M}}{\frac{1+c}{1-c}-\frac{1}{M-1}}\right)^{-\frac{1}{2}}\times \\ & \frac{1}{\sqrt{N}}\left[\left(\frac{\frac{1+c}{1-c}-\frac{1}{M}}{\frac{1+c}{1-c}-\frac{1}{M-1}}\right)^{-M\left(\frac{1+c}{1-c}\right)+1}\left(\frac{1+c}{1-c}-\frac{1}{M-1}\right)^{-\frac{1+c}{1-c}}\right]^N \,. \end{aligned} \tag{S42}$$

Substituting Eq. (S42) into Eq. (S41), we find that a single root exists for $M$ that gives the mode number of mating types that is straightforward to compute numerically with a standard root finding algorithm.

#### S5 Establishment probabilities

In the following we compute the establishment probability of a novel mating type in the population. We consider the case of facultative sex, i.e. the probability for asexual reproduction is given by $c \in (0, 1]$. Since the internal fixed point is stable in this case, see for example [IS87, Mating kinetics I], the calculation of the survival probability of the invading type gives a reasonable approximation of the establishment probability. Initially, when the mutant is still rare, the mutant individuals evolve independently. Hence, the mutant dynamics can be described by a branching process. For a general introduction on branching processes we refer to [HJV05, All11].

The birth and death probability for a rare invading mating type in a population of $M$ resident mating types can be written in terms of the transition probabilities given in Equation (S1). Assuming that the novel mating type has the index $M + 1$ and is present in $k$ copies we find that it increases by one with rate

$$T_k^+ = \sum_{j=1}^{M} \mathcal{T}_{(M+1)j} = \sum_{j=1}^{M} \left( c\frac{k}{N} + \frac{(1-c)}{2} \frac{k}{N} \frac{\sum_{i=1}^{M} n_i}{N} \right) \left( \frac{n_j}{N} \right) , \tag{S43}$$

and decreases by one with rate

$$T_k^- = \sum_{j=1}^{M} \mathcal{T}_{j(M+1)} = \sum_{j=1}^{M} \left( c\frac{n_j}{N} + \frac{(1-c)}{2} \frac{n_j}{N} \frac{\sum_{i \neq j} n_i}{N} \right) \left( \frac{k}{N} \right) . \tag{S44}$$

Assuming that the resident mating types are in the stationary state, i.e. setting $n_j = \frac{N}{M}$ for all resident mating types and therefore having $\sum_{j=1}^{M} n_j/N = 1$ we find

$$T_k^+ \approx c\frac{k}{N} + \frac{1-c}{2}\frac{k}{N} = \frac{k}{N}\frac{(1+c)}{2} \tag{S45}$$

and

$$\begin{aligned} T_k^- &\approx c\frac{k}{N} + \frac{k}{N}\frac{(1-c)}{2} \sum_{j=1}^{M} \frac{n_j}{N}\frac{N-n_j}{N} \\ &\approx c\frac{k}{N} + \frac{k}{N}\frac{(1-c)}{2}\left(1 - \frac{1}{M}\right) \\ &= \frac{k}{N}\frac{(1+c)}{2}\left(1-\frac{1}{M}\right) + \frac{k}{N}\frac{c}{M}. \end{aligned} \tag{S46}$$

Using the formula for the probability of extinction in a birth-death process (see e.g. [All11, Theorem 6.2])

$$p_{\text{ext}} = \frac{\sum_{k=1}^{\infty} \frac{T_1^- \cdots T_k^-}{T_1^+ \cdots T_k^+}}{1 + \sum_{k=1}^{\infty} \frac{T_1^- \cdots T_k^-}{T_1^+ \cdots T_k^+}}, \tag{S47}$$

we see that it depends on the ratio of the rates given in equations (S45) and (S46). We obtain

$$\frac{T_k^-}{T_k^+} = \frac{\frac{k}{2}(1+c)\left(1-\frac{1}{M}\right)+\frac{kc}{M}}{\frac{k}{2}(1+c)} = \left(1-\frac{1}{M}\right) + \frac{2c}{M(1+c)} = 1 - \frac{1-c}{M(1+c)}. \tag{S48}$$

Then equation (S47), using the geometric series, simplifies to

$$p_{\text{ext}} = \frac{\frac{1}{1-\left(1-\frac{1-c}{M(1+c)}\right)} - 1}{1 + \left(\frac{1}{1-(1-\frac{1-c}{M(1+c)})} - 1\right)} = 1 - \frac{1-c}{M(1+c)}. \tag{S49}$$

This results in a survival probability of

$$p_{\text{surv}} = 1 - p_{\text{ext}} = \frac{1}{M}\frac{(1-c)}{(1+c)}. \tag{S50}$$

Since $Q_M^{\text{Est}}$ equals the survival probability of this process we have found the expression in Eq. (13) from the Main Text.

We note that for $c = 1$, i.e. exclusively clonal reproduction, our model reduces to a multi-allelic Moran model. In this setting the establishment probabilities can not be calculated via the here implemented approach.

### S6 Breakdown of the mean extinction time calculation

In the main text, we calculate an approximation for the mean extinction time as a function of the population size, $N$, rate of asexual reproduction $c$ and number of resident mating types, $M$ (see Eq. (18)). We also show that in certain regions of parameter space, this approximation breaks down (see Eq. (20)). In these parameter regions, in which genetic drift dominates the dynamics, the mean time to extinction of a resident allele is better approximated by the neutral multi-allelic Moran model (see below).

In the following we show that it is the inaccuracies of our quasi-stationary linear approximation for the distribution of mating type allele frequencies about deterministic fixed points (see Section S2) that drives this break down. This approximation assumes that a given fixed point of the deterministic system is sufficiently stable that the distribution of allele frequencies around this fixed point can be well captured by a Gaussian (normal) distribution (see Eq. (S25)). Clearly this local description of the distribution does not account for the behaviour of the system at the extinction boundaries. When population sizes are high, rates of asexual reproduction low, and the number of resident mating types small, the probability mass predicted by the Gaussian approximation at these extinction boundaries is negligible. In this regime our approximation continues to be accurate, as illustrated in Figure S1. However, outside this range (i.e when $N$ is low, $c$ high, or $M$ large), the variance of the Gaussian approximation becomes sufficiently large that non-negligible probability mass is predicted at the boundary (see Figure S3). These parameter regions correspond to areas where drift dominates the dynamics.

In regions of parameter space where the Gaussian approximation becomes inaccurate, we expect our expression for the distribution of mating types, $\mathcal{P}_M^{\text{st}}$, (which relies on the Gaussian approximation, see Eq. (S28) and the subsequent calculations) to also become inaccurate. However, as these inaccuracies occur in biologically less interesting regions of parameter space where mating types are frequently becoming extinct, they do not affect the dominant modes of $\mathcal{P}_M^{\text{st}}$ (see Figure 1 in the Main Text). In contrast, in our investigation of extinction times we are sometimes probing very unstable configurations of the resident mating types (see Figure 5 in the Main Text), and thus the approximation breaks down in these parameter regions.

We now show that it is indeed the Gaussian approximation for the quasi-

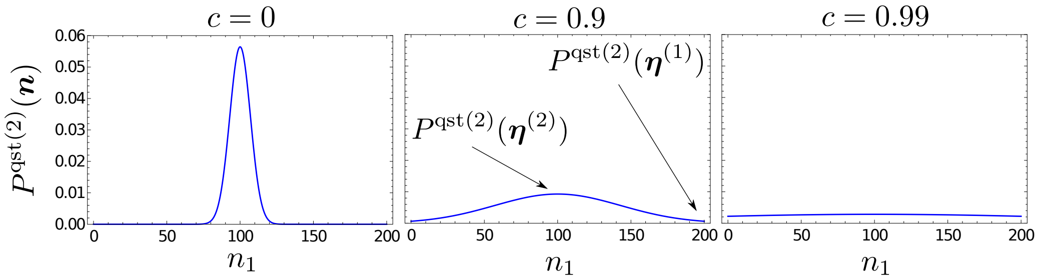


Figure S3: Figure illustrating the breakdown in the quasi-stationary approximation for the distribution of alleles around a fixed point, $P^{\mathrm{qst}}(\boldsymbol{n})$ (see Eq. (S27)). We consider two mating types in a fairly small population, $M = 2$, $N = 200$. For $c = 0$, the approximation remains physically reasonable. As $c$ is increased however, a non-negligible amount of probability mass builds up on the boundary relative to that at the fixed point around which the distribution is centered (that is, the ratio of $P^{\mathrm{qst}(2)}(\boldsymbol{\eta}^{(1)})$ to $P^{\mathrm{qst}(2)}(\boldsymbol{\eta}^{(2)})$ tends to one). A similar pattern can be observed as $N$ decreases and $M$ increases.

stationary distributions of mating type allele frequencies about deterministic fixed points that drives the breakdown of the extinction time calculation. In Figure S4 we plot the probability mass predicted by the Gaussian distribution at an extinction boundary relative to the distribution's value at the corresponding fixed point. This quantity should be very low for the approximation to remain physically reasonable (i.e. there should be very little mass at the extinction boundary), while it approaches one as the predicted distribution becomes increasingly flat and inaccurate. We observe that high values of predicted probability mass at the extinction boundary coincide with regions where our prediction for the mean time to extinction, Eq. (18), break down, and the neutral theory becomes more appropriate (see Eq. (20)).

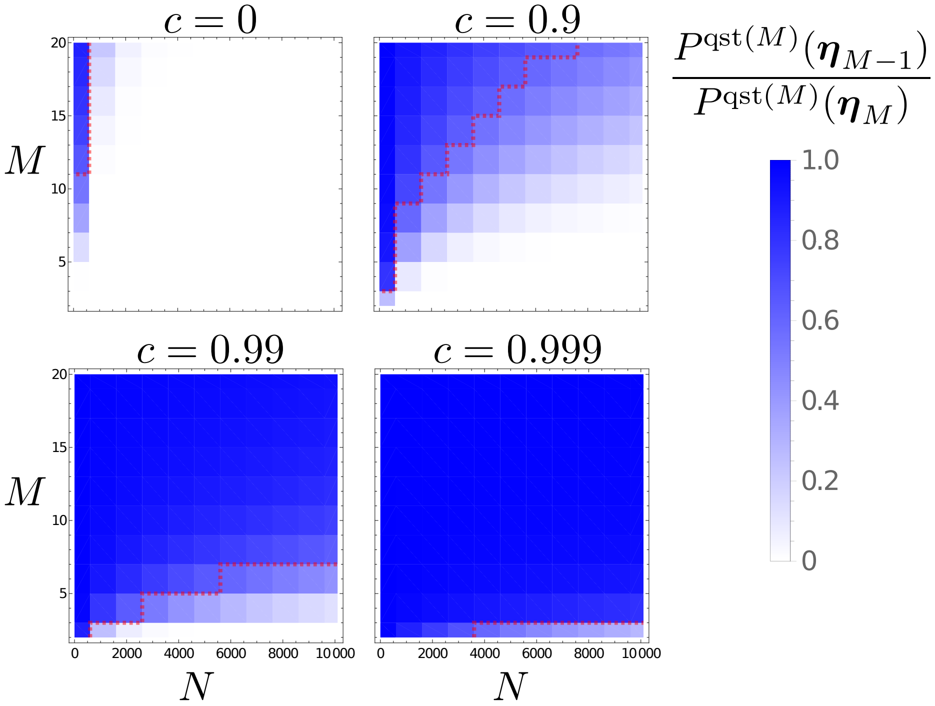


Figure S4: Figure illustrating the coincidence of the breakdown in the quasi-stationary approximation for the distribution of alleles around a fixed point, $P^{\mathrm{qst}}(\boldsymbol{n})$ (see Eq. (S27)). The colorbar indicates the probability mass predicted by the quasi-stationary distribution at the nearest extinction boundary, relative to the probability mass at the center of the quasi-stationary distribution (i.e. at the deterministic fixed point). Low values (in white) are associated with regions of parameter space where the quasi-stationary distribution remains reasonable. High values (in deep blue) are associated with regions where the quasi-stationary distribution becomes a poor approximation (see Figure S3). Areas to the top left of the red dashed lines are those where the conditions in Eq. (18) are violated (that is, our approximation for the mean time to fixation breaks down).

#### S7 Neutral mean extinction time

We derive an approximation for the neutral mean extinction time as given in Eq. (19) in the Main Text. Therefore, we identify our process with the multi-allelic neutral Moran model. This model has been analysed in detail in [BBM07]. Using their Eq. (44) and plugging in our parameter values, i.e. $\boldsymbol{x}(0) = \boldsymbol{x}^* = 1/M$ and $r = 1$ we find

$$\tau = -N \sum_{s=1}^{M-1} (-1)^{s-1} \binom{M}{s} \frac{s}{M} \log\left(\frac{s}{M}\right), \tag{S51}$$

which is the result in Eq. (19) from the Main Text. Note, that there is a time-scale difference of 1/2 between their model (Wright-Fisher diffusion) and our implementation (Moran model) explaining. Furthermore, we consider the dynamics on the original time-scale resulting in the factor $N$ in front of the sum (see also their comment preceding their Eq. (2)).

#### S8 The stationary distribution $\mathcal{P}_M^{\text{st}}$ for the parameters given in Table 1.

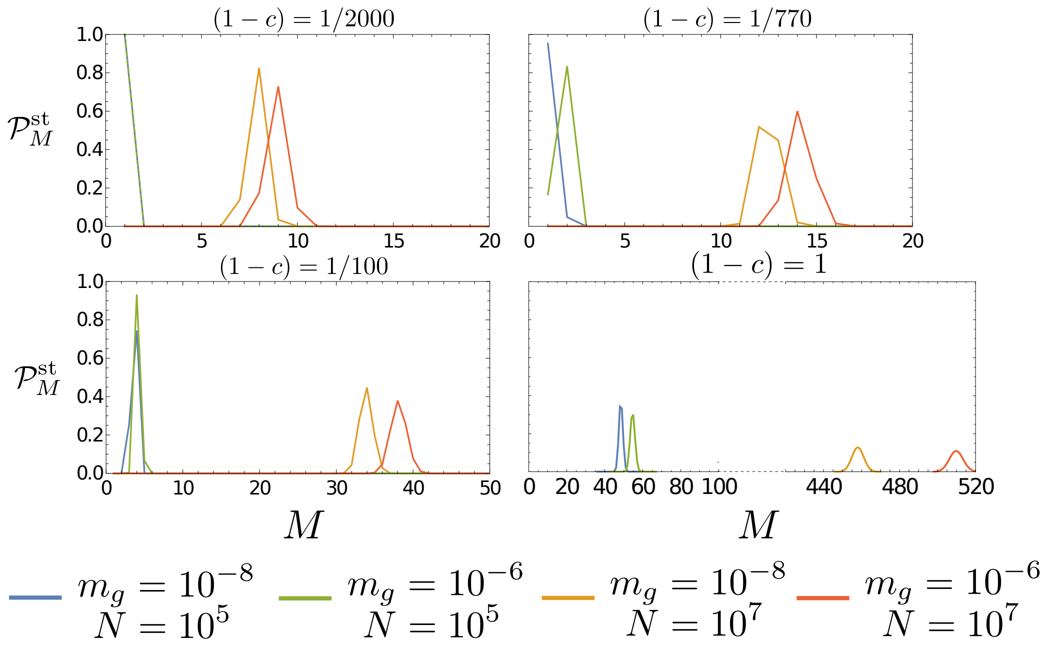


Figure S5: Theoretical distributions for $\mathcal{P}_M^{\text{st}}$ (see Eq. (9) in the Main Text) using parameter values from Table 1 in the Main Text. Panel top-left ($(1-c) = 1/2000$): parameters estimated for *S. cerevisiae*. Panel top-right ($(1-c) = 1/770$): parameters estimated for *C. reinhardtii*. Panel bottom-left ($(1-c) = 1/100$): parameters estimated for Tetrahymena. Panel bottom-right ($(1-c) = 1$): parameters estimated for *S. commune*. Note that for clarity the scale of the $M$ axis changes between panels. Also note that for the final panel (bottom-right) the region $100 \leq M \leq 400$ has been omitted.

### References


[All11] Linda J.S. Allen. *An Introduction to Stochastic Processes with Applications to Biology.* Pearson Education, New Jersey, 2nd edition, 2011.

[AS65] M. Abramowitz and I. A. Stegun, editors. *Handbook of Mathematical Functions: with Formulas, Graphs, and Mathematical Tables.* Dover Publications, New York, 1965.

[BBM07] Gareth J. Baxter, Richard A. Blythe, and Alan J. McKane. Exact solution of the multi-allelic diffusion model. *Mathematical biosciences*, 209 1:124–70, 2007.

[CK18] G.W.A Constable and H. Kokko. The rate of facultative sex governs the number of expected mating types in isogamous species. *Nat. Ecol. Evol.*, 2(7):1168–1175, 2018.

[EK86] Stewart N. Ethier and Thomas G. Kurtz. *Markov processes: characterization and convergence.* Wiley series in probability and mathematical statistics. J. Wiley & Sons, New York, Chichester, 1986.

[HJ91] R. A. Horn and C. R. Johnson. *Topics in Matrix Analysis.* Cambridge University Press, Cambridge, 1991.

[HJV05] Patsy Haccou, Peter Jagers, and Vladimir A. Vatutin. *Branching Processes: Variation, Growth, and Extinction of Populations.* Cambridge Studies in Adaptive Dynamics. Cambridge University Press, 2005.

[IS87] Y. Iwasa and A. Sasaki. Evolution of the number of sexes. *Evolution*, 41(1):49–65, 1987.

[Kam07] NG Van Kampen. *Stochastic processes in physics and chemistry.* North Holland, 2007.

[MBR14] A. J. McKane, T. Biancalani, and T. Rogers. Stochastic pattern formation and spontaneous polarisation: the linear noise approximation and beyond. *Bull. Math. Biol.*, 76:895–921, 2014.
